# Single cell genomic characterization reveals the cellular reprogramming of the gastric tumor microenvironment

**DOI:** 10.1101/783027

**Authors:** Anuja Sathe, Sue Grimes, Billy T. Lau, Jiamin Chen, Carlos Suarez, Robert Huang, George Poultsides, Hanlee P. Ji

**Author notes:** Corresponding author: Hanlee P. Ji.

## Abstract

**Purpose:** The tumor microenvironment **(TME)** consists of a heterogenous cellular milieu that can influence cancer cell behavior. The characteristics of the cellular TME have a dramatic impact on treatments such as immunotherapy. These features can be revealed with single-cell RNA sequencing **(scRNA-seq)**. We hypothesized that single cell gene expression studies of gastric cancer **(GC)** together with paired normal tissue and peripheral blood mononuclear cells **(PBMCs)** would identify critical elements of cellular dysregulation not apparent with other approaches.

**Methods:** Single cell gene expression studies were conducted on seven patients with GC and one patient with intestinal metaplasia. We sequenced 56,167 cells comprising GC (32,407 cells), paired normal tissue (18,657 cells) and PBMCs (5,103 cells). Protein expression of genes of interest was validated by multiplex immunofluorescence.

**Results:** Tumor epithelium had copy number alterations and a distinct gene expression program compared to normal with intra-tumor heterogeneity. The GC TME was significantly enriched for stromal cells, macrophages, dendritic cells **(DCs)** and Tregs. TME-exclusive stromal cells expressed extracellular matrix components distinct from normal tissue. Macrophages were transcriptionally heterogenous and did not conform to a binary M1/M2 paradigm. Gene expression program of tumor DCs was unique from PBMC DCs. TME-specific cytotoxic T cells comprised of two exhausted heterogenous subsets. Helper, cytotoxic T, Treg and NK cells expressed multiple immune checkpoint or costimulatory molecules. Receptor-ligand analysis revealed TME-exclusive inter-cellular communication.

**Conclusions:** Single cell gene expression studies revealed widespread reprogramming across multiple cellular elements in the milieu of the GC TME. Cellular remodeling was delineated by changes in cell numbers, transcriptional states and inter-cellular interactions. This characterization facilitates understanding of tumor biology and enables the identification of novel molecular targets including for cancer immunotherapy.

**STATEMENT OF TRANSLATIONAL RELEVANCE:** We leveraged the power of single-cell genomics to characterize the heterogenous cell types and states in the tumor microenvironment (TME). By profiling thousands of single cells from surgical resections of gastric cancer together with paired normal mucosa and peripheral blood mononuclear cells (PBMCs), we determined the deviations in the TME from physiological conditions. Our analysis revealed a cellular reprogramming of the TME compared to normal mucosa in immune and stromal lineages. We detected transcriptional heterogeneity within macrophages and a TME-specific gene expression program in dendritic cells. Cytotoxic T cells in the TME had heterogenous profiles of exhaustion and expression of multiple immune checkpoint and costimulatory molecules. We constructed a receptor-ligand based inter-cellular communications network that was exclusive to tumor tissue. These discoveries provide information at a highly granular cellular resolution enabling advances in cancer biology, biomarker discovery and identification of treatment targets such as for immunotherapy.

## INTRODUCTION

Gastric cancer **(GC)** is the fifth most common cancer and the third leading cause of cancer deaths worldwide (1). The current histopathologic classification scheme designates GCs as either intestinal or diffuse subtype according to the differentiation and cohesiveness of glandular cells and other features of cellular morphology (2). Intestinal GC is preceded by changes in the gastric mucosa called the Correa cascade that progresses through inflammation, metaplasia, dysplasia and adenocarcinoma. Diffuse GCs lack intercellular adhesion and exhibit a diffuse invasive growth pattern. Recent integrated genomic and proteomic analyses including by the Cancer Genome Atlas **(TCGA)** and the Asian Cancer Research Group **(ACRG)** (4, 5) have refined the classification of GC and revealed that they fall into several distinct molecular subtypes that include the intestinal and diffuse classification. Regardless of the histopathologic or molecular subtype, GCs are not isolated masses of cancer epithelial cells – rather they have a complex morphology where cancer cells are embedded and/or surrounded by a diverse assortment of different cell types (3). Referred to as the tumor microenvironment **(TME)**, this cellular milieu contains fibroblasts, endothelial, immune and other diverse cell types.

Interactions among the cellular components of the TME and cancer cells enable tumors to proliferate, metastasize and alter the response of the immune system. Increasingly, it is recognized that the cellular features and interactions of the TME components play an important role in GC biology and determine the efficacy of specific treatments such as immunotherapy. There are general mechanisms where gastric tumor cells suppress the local immune cells and dysregulate the patient’s immune system (4). Moreover, tumors are stimulated by various growth factors that originate from the cellular compartments within the TME. Thus, the cellular characterization of the TME provides a more sophisticated picture of the context of tumor cell growth within its tissue of origin, characteristics of immune infiltrate and inter-cellular interactions. These are important elements that determine therapeutic response but have not been characterized.

The major objective of this study was to determine the specific cellular and transcriptional features that distinguish the GC TME from normal gastric tissue. We sought to define these differences at the resolution of single cells with single cell RNA-seq **(scRNA-seq)**. We delineated cell-specific features that are otherwise lost when using “bulk” methods in which molecular analytes cannot be attributed to their cell-of-origin. Our study used matched gastric tumor and normal tissue from the same patients. We made comparisons among these paired normal-tumor samples as well as extrapolated the more global cellular TME differences noted among all of the samples. In addition, we had peripheral blood mononuclear cells **(PBMCs)** for a subset of patients from which we made comparisons with tumor infiltrating immune cells. Our analysis considered both the pathologic category as well as distinguishing features that delineate the molecular subtype.

To identify the cellular and transcriptional features of cells in the gastric TME, we developed an extensive analytical framework (**Figure 1A**) (5–8). With single cell resolution, we identified multiple differences between the tumor and normal epithelium and conducted differential gene expression analysis for each cell type in the context of the gastric TME versus the normal stomach. Our results indicated specific cellular programming states based on changes seen in gene expression and extrapolated cellular regulatory networks. Finally, we identified important inter-cellular communication networks specific to the gastric TME compared to the gastric normal tissue. Our study identified cellular and biological features that are specific to the TME, and thus offer insight which may help infer new therapeutic targets.

**Figure 1:**
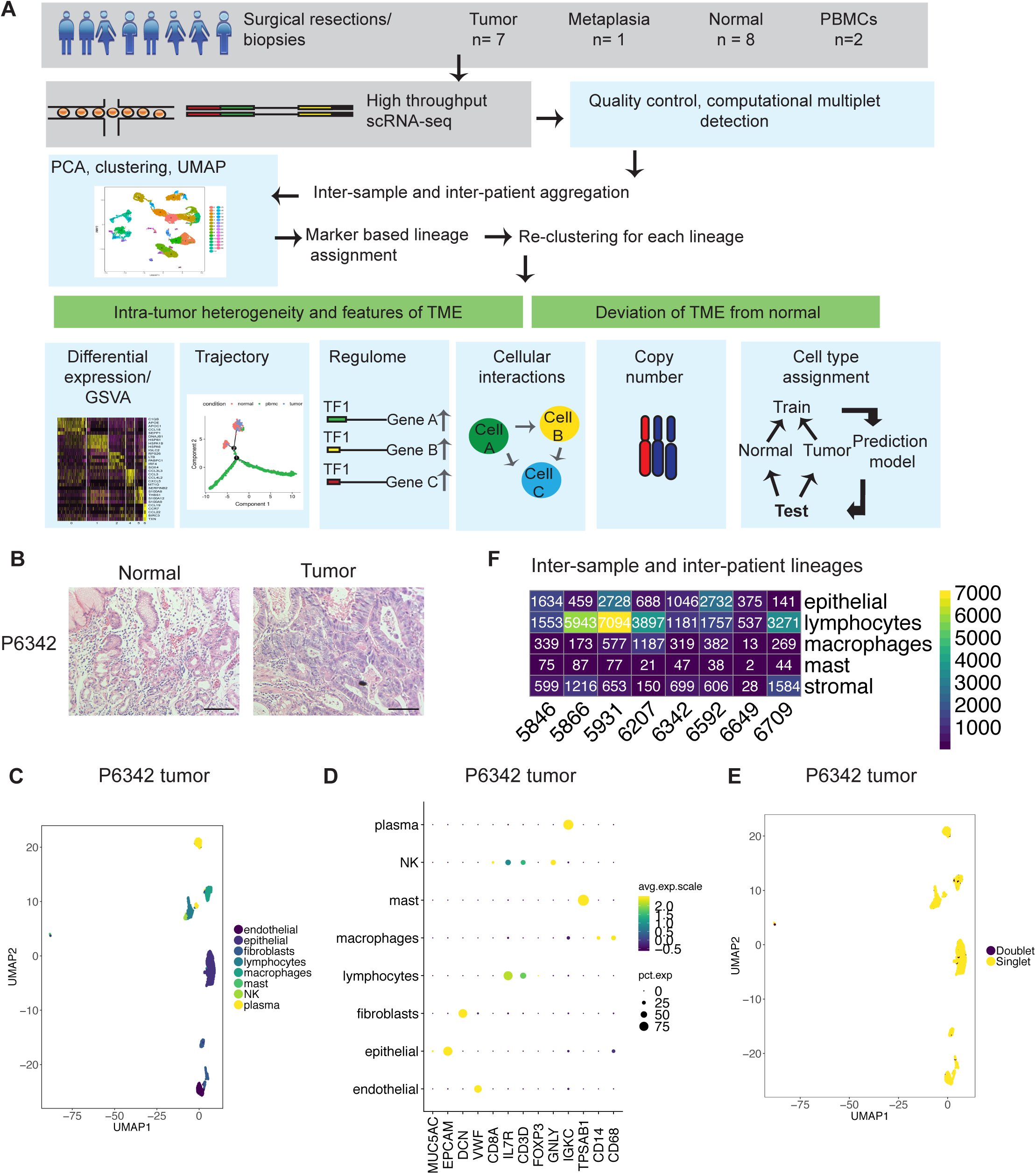
(A) Schematic representation of experimental design and analytical methods used in this study. (B) Representative images of hematoxylin and eosin staining of FFPE tissue from P6342. Scale bar indicates 50 μm. (C-F) Example of clustering analysis in tumor sample of P6342. (C) UMAP representation of dimensionally reduced data following graph-based clustering with marker-based cell type assignments. (D) Dot plot depicting expression levels of specific lineage-based marker genes together with the percentage of cells expressing the marker. (E) UMAP representation of dimensionally reduced data following graph-based clustering with computational doublet identification. (F) Heatmap depicting number of cells identified in aggregated analysis for each lineage per patient.

## METHODS

### Sample acquisition and tissue processing

All samples were acquired with informed consent under an approved institutional review board protocol from Stanford University. Gastric tumor and metaplasia samples were obtained from surgical resections or endoscopic biopsies. In parallel, we also obtained tissue from matched normal gastric sites that were displaced at least several centimeters from the tumor. The normal gastric tissue was confirmed to lack tumor cells based on histopathology review.

Tissues were collected in plain RPMI immediately after surgical resection and stored on ice. Following dissection with iris scissors, tissue fragments were subjected to enzymatic and mechanical dissociation using human tumor dissociation kit (Miltentyi Biotec, Germany) with the Gentle MACS Octodissociator (Miltenyi) as per manufacturer’s protocol with the ‘37_h_TDK_3’ program. For biopsy specimens, half the volume of media and enzymes were used.

Dissociated cells were incubated with RBC lysis buffer (155 mM ammonium chloride, 10 mM potassium bicarbonate, 0.1 mM EDTA) for 10 minutes followed by neutralization with PBS. All centrifugation steps were carried out at 400g for 3 minutes.

Blood collected in EDTA or sodium heparin was overlaid on 15 ml Ficoll-Paque Plus (GE healthcare, Chicago, IL) contained in a Sepmate 50 ml tube (Stemcell Technologies, Vancouver, Canada), and centrifuged for 10 minutes at 1200g. Interphase containing the PBMCs was decanted into a fresh tube followed by 2 washes with PBS with centrifugation at 400g for 3 minutes. Cells were cryofrozen using 10% DMSO in 90% FBS (Thermofisher Scientific, Waltham, MA) or Cryostor CS10 (StemCell Technologies) (for P6709 samples) in a CoolCell freezing container (Larkspur, CA) at −80 °C for 48 hours followed by storage in liquid nitrogen.

Dissociated cells or PBMCs were washed twice in RPMI + 10% FBS, filtered through 70 μm (Flowmi, Bel-Art SP Scienceware, Wayne, NJ), followed by 30 μm (Miltenyi) or 40 μm filter (Flowmi). Cryofrozen cells were rapidly thawed in a bead bath at 37 °C followed by the above steps. Live cell counts were obtained on a BioRad TC20 cell counter (Biorad, Hercules, CA) using 1:1 trypan blue dilution. Cells were concentrated between 500-1500 live cells/μl and used for loading single-cell reactions.

### Single cell sequencing

Samples from each patient were processed in one batch for library preparation **(Supplementary Table S1**). Chromium Single Cell 3′ Library & Gel Bead Kit v2 (10x Genomics, Pleasanton, CA, USA) was used as per manufacturer’s protocol. For a given sample, 10000 cells were targeted for tissue dissociation suspensions and 3000 for PBMCs with 14 PCR cycles for cDNA and library amplification using a program as listed in the manufacturer’s protocol. A 1% or 2% E-Gel (Thermofisher Scientific) were used for quality control evaluation of the amplified cDNA and sequencing libraries. A Qubit (Thermofisher Scientific) or qPCR with Kapa library quantification kit (Kapa Biosystems, Wilmington, MA) was used to quantify the libraries as per the manufacturer’s protocol. We conducted sequencing on an Illumina sequencer (Illumina, San Diego, CA) **(Supplementary Table S1**). Cell Ranger v 3.0 (10x Genomics) ‘mkfastq’ and ‘count’ commands were used with default parameters and alignment to GRCh38 to generate matrix of unique molecular identifier **(UMI)** counts per gene and associated cell barcode. Datasets are available under dbGAP identifier phs001818.v1.p1.

### Clustering analysis

We used Seurat (v2.3.4) (8) to create data objects from the matrix outputs. We removed cells that expressed fewer than 200 genes, had greater than 20% mitochondrial genes or had number of UMI in an outlier range indicative of potential doublets **(Supplementary Table S1**). We excluded genes detected in fewer than three cells. Data was normalized to log scale using the ‘NormalizeData’ function with a default scale parameter of 10000. Highly variable genes were identified using the ‘FindVariableGenes’ function with parameters for x.low.cutoff=0.0125, x.high.cutoff=6 and y.cutoff=0.5. The effects of variation in sequencing depth were regressed out by including ‘nUMI’ as a parameter in the ‘ScaleData’ function. These variable genes were used as input for PCA using the ‘RunPCA’ function. The first 20 principal components **(PCs)** and a resolution of 0.8 were used for clustering using ‘FindClusters’. UMAP was used for two-dimensional representation of first 20 PCs with ‘RunUMAP’.

Differential gene expression for identifying markers of a cluster relative to all other clusters or compared to a specific cluster was determined using the ‘FindAllMarkers’ or ‘FindMarkers’ functions respectively. Parameters provided for these functions were: genes detected in at least 25% cells; differential expression threshold of 0.25 log fold change using Wilcoxon rank sum test with p < 0.05 following Bonferroni correction. We compared the marker genes for each cluster to literature-based markers of cell lineages to assign a cell lineage per cluster **(Supplementary Table S2**).

Individual Seurat data objects were merged iteratively using the ‘MergeSeurat’ function after filtering doublets identified by DoubletFinder, an R package that enables computational identification of doublets (**Supplementary methods**). The merged object was processed as described above with library preparation batch and number of UMIs **(Supplementary Table S1**) included as parameters for regression in the ‘ScaleData’ function to regress batch effects and variation in sequencing depth respectively. The ‘DoHeatmap’, ‘FeaturePlot’, ‘DimPlot’, DotPlot’, ‘VlnPlot’ were used for visualization.

For a secondary cluster analysis of each cell lineage from this aggregated dataset, clusters of interest were identified and subset using ‘SubsetData’ with parameter ‘do.clean’ set to true. Detection of variable genes, scaling with UMI regression, PCA, clustering and UMAP were repeated as described above. Following this step, we removed clusters with co-expression of cell lineage markers as multiplets **(Supplementary Table S3**). Proportions of each cell type relative to the total number of cells in the sample were compared for tumor and normal sites using two proportions z-test and represented as box plots after re-clustering analysis for each lineage. Additional description of methods is available under **Supplementary Methods**.

## RESULTS

### Cohort of gastric cancer and intestinal metaplasia

We obtained tissues from surgical resections or endoscopic biopsies from seven patients with GC and one patient with gastrointestinal metaplasia **(GIM)**. These tissue samples represented paired gastric tumor and gastric normal tissue from the same patient derived from the same anatomical region of the stomach. Specifically, four tumors were located at the gastroesophageal junction **(GEJ)** and four were located in the body and antrum. Also, we analyzed matched peripheral blood mononuclear cells **(PBMCs)** from two patients. Based on histopathology review, the GC tumors had intestinal, diffuse or mixed features **(SupplementaryTable S4**, **Figure 1B**, **Supplementary Figure 1A**). The majority of samples had immunohistochemistry **(IHC)** testing for protein expression of MLH1, MSH2, MSH6 and PMS2 which are proteins involved in DNA mismatch repair. We used these IHC results to classify tumors into microsatellite stability **(MSS)** or microsatellite instability **(MSI)** molecular subtypes **(Supplementary Table S4, Supplementary Methods)**. Histopathology showed that three patients (P5866, P6207, P6342) had active gastritis or intestinal metaplasia in paired non-malignant tissue.

### Single-cell transcriptomic profiles from gastric cancer

With scRNA-seq, we obtained transcriptional profiles of 32,407 single cells from tumors or metaplasia, 18,657 single cells from paired normal tissue and 5,103 PBMCs **(Supplementary Table S1**). To determine cell type and gene expression changes, we employed a series of analytical steps for each individual scRNA-seq dataset. This included quality filtering the single cell data **(Methods)**, principal component analysis **(PCA)** on genes that were variably expressed across cells and graph-based clustering on the first twenty principal components (8, 9). In addition, we employed uniform manifold approximation and projection **(UMAP)** to reduce the dimensionality of this data and allow the visualization of cell-type clusters defined by their transcriptional profiles.

Differentially expressed **(DE)** genes were identified by selecting genes expressed in greater than 25% of cells in a cluster and having a log fold change greater than 0.25, using a cut-off of p < 0.05 following Bonferroni correction. We compared the DE genes from each cluster to known marker genes of various cell types **(Supplementary Table S2**). This information enabled us to link clusters to specific cell types including epithelial cells (expressing *PGC*, *TFF1*, *MUC5AC*, *EPCAM*, *GIF*, *CHGA*), fibroblasts (*THY1*, *DCN*, *COL4A1*, *FAP*), endothelial cells (*PECAM*, *ENG*, *VWF*, *SELE*), immune cells (*PTPRC*) such as CD4 T (*CD3D*, *CD4*, *IL7R*), cytotoxic T (*CD3D*, *CD8A*, *CD8B*), regulatory T (*FOXP3*, *IL2RA*), NK (*NKG7*, *GNLY*), B (*MS4A1*), plasma cells (immunoglobulin genes), mast cells (*TPSAB1*) and macrophages (*CD68*, *CD14*, *FCGR3A*). Examination of cells expressing markers of disparate cell types facilitated the computational detection of doublets (**Figure 1C, 1D**, **Supplementary Methods)**.

### Data integration for joint cell analysis across all samples

To determine both the similarities and differences among all of the matched samples including GC, metaplasia and normal gastric cells, we aggregated all of the scRNA-seq data into a single data matrix. This integrated data set combined the tumor, metaplastic, normal sites and PBMCs scRNA-Seq results across all samples and patients. To reduce experimental variance for this combined data set, we regressed out the batch effects from library preparation and variation in sequencing depth for all of the samples and also removed computationally detected doublets **(Methods, Supplementary Methods)**.

Overall, PCA clustering and UMAP analysis identified 40 distinct cellular clusters that were specific for a variety of cell types including epithelial, stromal (fibroblasts, endothelial cells), lymphocytes, macrophages and mast cells – these cell types were found among all of the patient tissue samples **(Supplementary Figures 1B and 1C, Figure 1E**). On closer examination, each cell type had multiple and distinct transcriptional states. For additional characterization of transcriptional states across all patients, we aggregated the data for each cell type across samples and conducted a clustering analysis **(Methods)**.

### Classifying tumor, normal and metaplastic epithelial cell populations

We detected differences between tumor, metaplastic and normal epithelial cells, differences in tumor epithelium derived from different patients as well as intra-tumoral sub-clonal heterogeneity within an individual tumor.

We identified epithelial cells which expressed *PGC*, *TFF1*, *MUC5AC*, *EPCAM*, *GIF*, *CHGA* (**Supplementary Figure 2A, 2B)** from our integrated dataset. On close inspection, we identified three subclasses of epithelial cells. The first subclass consisted of normal gastric epithelial cells – over 80% of these cells came from the normal gastric tissue samples (**Figure 2A, B, C**). All samples had normal epithelial cells regardless of whether these cells originated from gastric tumor, normal or metaplasia tissue (**Supplementary Figure 2C**).

**Figure 2:**
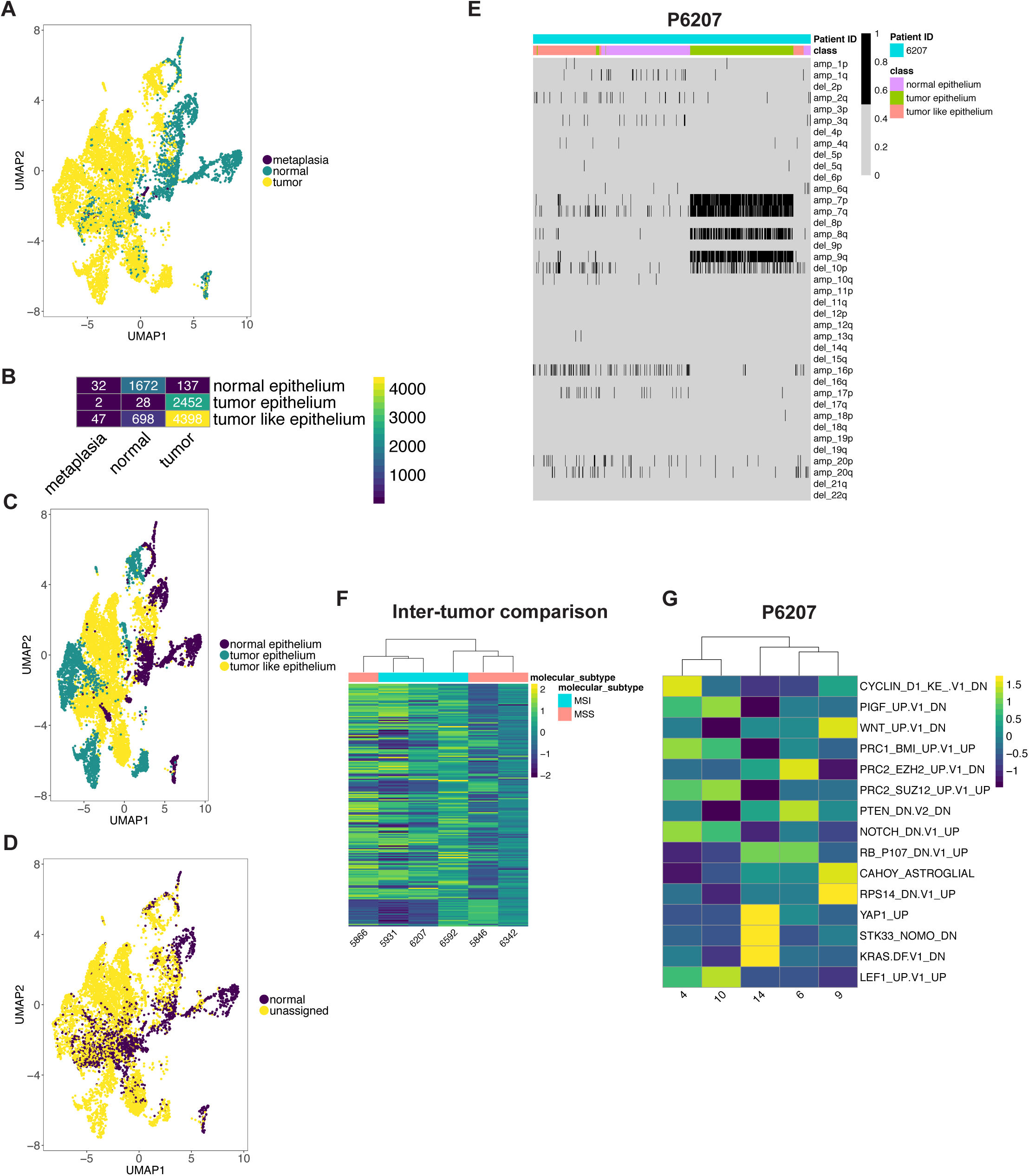
(A) UMAP representation of epithelial cells following graph-based clustering colored by sample origin. (B) Heatmap depicting number of cells per defined epithelial class according the sample origin. (C-D) UMAP representation of epithelial cells following graph-based clustering colored by (C) class (D) predicted class according to scPred. (E) Heatmap representation of statistically significant copy number changes for depicted chromosomes for epithelial cells from P6207 as a representative example. ‘amp’ denotes amplification, ‘del’ denotes deletion. (F-G) Heatmaps depicting average gene set activity of top MSigDB oncogenic c6 gene signatures following GSVA (ANOVA FDR p value < 0.05) across tumor epithelial clusters for (F) all patients and (G) all clusters for P6207.

The second subclass consisted of tumor-specific epithelial cells – with 98% of these cells originating from the gastric carcinoma samples. Interestingly, each cluster in this subclass were dominated by a single gastric tumor sample (**Supplementary Figure 2C**) – this result indicated the extent of inter-tumor heterogeneity among all of the GCs, meaning that each individual tumor had distinct transcriptional properties.

The third subclass involved epithelial cells which were tumor-like. This class contained approximately 50% of cells from a patient P6649 who had gastric metaplasia but did not have GC. Given their origin and histopathologic result, we classified these cells as representing metaplastic or dysplastic epithelial cells, which had a subset of transcriptional features that overlapped with the tumor epithelial cell sub-category.

As an independent determination of gastric tumor versus normal epithelial cells, we employed a completely different method that uses a supervised machine learning algorithm. Called scPred, this approach eliminates the statistical inconsistencies seen with other single cell methods and thus provides highly accurate cell type assignment (10). We applied scPred on gastric epithelial cells derived from normal and tumor tissue. Specifically, we randomly subset tumor and normal site derived cells into two. One subset from each of these classes was included in the training dataset to build the scPred prediction model. We tested this model on all of the remaining cells for all samples. When we compared the scPred results to our Seurat analysis, the results were concordant. For example, seventy-seven percent of cells in the tumor-like epithelium class were found to be distinct and not classified as normal gastric epithelial cells (**Figure 2D**, **Supplementary Figure 2D**). Thus, scPred confirmed the specific transcriptional differences among the normal, tumor and tumor-like epithelial subclasses.

### Gene expression differences among the epithelial cell subclasses

Specific gene expression differences distinguished the three subclasses of epithelial cells. Our analysis of the normal gastric epithelium identified distinct mucosal cell populations such as pit cells, mucous neck cells, zymogen secreting chief cells, intrinsic factor producing parietal cells and neuroendocrine cells (**Supplementary Figure 2B**). In contrast, tumor epithelium and tumor-like epithelium downregulated some of these gastric mucosa marker genes such as *MUC6*, *TFF2*, *TFF1*, *MUC5AC* **(Supplementary Table S5**). Tumor epithelium had increased expression of intestinal mucosa markers *TFF3, FABP1, SPINK4, MUC13* and *REG4*. Tumor-like epithelium had significantly increased expression of the previously identified gastric cancer marker genes *KRT7* and *KRT17* (11), *SOX4* and *HES1* that have been implicated in metaplasia pathogenesis (12). Compared to normal gastric cells, both tumor and tumor-like epithelial cells had upregulation of gene sets that included pathways for Myc, DNA repair and Notch signaling (**Supplementary Figure 2E**). However, only tumor cells had higher upregulation of EMT and KRAS signaling compared to the other epithelial cell classes.

### Copy number alterations distinguish tumor and normal epithelium

Using scRNA-seq, one can detect large copy number changes within individual cells (13). Specifically, DNA copy number changes such as amplifications or deletions over extended segments of the genome (megabases involving chromosome arms) leads to a concomitant increase or decrease in expression for genes in that segment **(Supplementary Methods)**. We determined which cells had these large copy number changes among our samples. Copy number changes were inferred according to the posterior probability for each cell to belong to one of the components with lower or higher gene expression indicative of deletion or amplification respectively. We excluded the sample from patient P6709 from this analysis since we only detected 21 epithelial cells from the tumor site, possibly indicative of response to neoadjuvant chemotherapy **(Supplementary Figures 2B, 1A, Supplementary table 4**).

To identify copy number alterations, we analyzed each patient’s normal epithelial cells versus the matched tumor or tumor-like epithelial cells (**Figure 2E**, **Supplementary Figure 3A**). A wide spectrum of chromosome arm imbalances were present in these tumors. For patient P6207, we detected genomic amplifications in chromosome arms 7p, 7q, 8q and 9q. Also, we identified a deletion in 10p. Patient P6342’s tumor had amplification of 20q with deletion of 4q. Patient P5846’s tumor had an amplification of 19q. Patient P5866’s tumor had deletion of 16q. Patient P5931’s tumor also had amplifications of 7p and 7q that have been previously described in gastric cancer (14). For Patient P6649 with metaplasia and P6592, samples contained only a small number of cells with significant copy number changes. Overall, these results provide additional orthogonal confirmation of the distinction between normal and tumor epithelium.

### Cancer cell differences across samples and tumor clonal heterogeneity

We analyzed tumor and tumor-like epithelial cells for the transcriptional activation of various oncogenic pathways **(Supplementary Methods).** We observed significant differences in activation levels across patients (ANOVA FDR p value < 0.05) indicating variation in cancer cells between patients (**Figure 2F**). Interestingly, these activation profiles did not cluster according to differences between the molecular subtypes of MSI and MSS.

The individual patient tumors contained multiple clusters of tumor or tumor-like cells, an indication of intra-tumoral sub-clonal heterogeneity. We conducted a pathway analysis to understand differences in signaling pathway activation across these subclones within a patient’s tumor. Upon hierarchical clustering, pathway activation profiles grouped into three to five sub-populations for each patient’s tumor confirming a sub-clonal structure (**Figure 2G**, **Supplementary Figure 4**). For example, heterogeneity in P6207 was characterized by differences in cell cycle, KRAS pathway, Wnt activation (**Figure 2G**). We postulate that these sub-populations may have a distinct growth advantage compared to other cells lacking these changes.

### TME reprogramming leads to macrophage states other than M1 and M2 classification

TME macrophages had distinct cellular and gene expression changes compared to normal gastric tissue, some of which have not been described. This result is consistent with the observation that macrophages acquire heterogenous phenotypes depending on their activating stimulus (15). Macrophage phenotypes are called M1 or M2 with anti and pro-tumorigenic functions respectively. However, the gene expression signatures we observed did not fall in line with either the canonical M1 or M2 classes.

First, we detected various subclasses of myeloid lineage cells as seen with 11 distinct clusters detected as determined by marker genes (**Figure 3A-D**). This was observed consistently across all patient samples (**Supplementary Figure 5A**). The nine monocyte-macrophage clusters were defined by marker gene expression of *CD14, FCGR3A, CD68*. The two dendritic cell **(DC)** clusters were defined by marker gene expression of *CLEC4C, ID2, IRF4, CD83* (**Figure 3C**). Notably, macrophages and DCs were significantly enriched in tumor compared to normal gastric tissue (z test of proportions, p < e-07) (**Figure 3E**).

**Figure 3:**
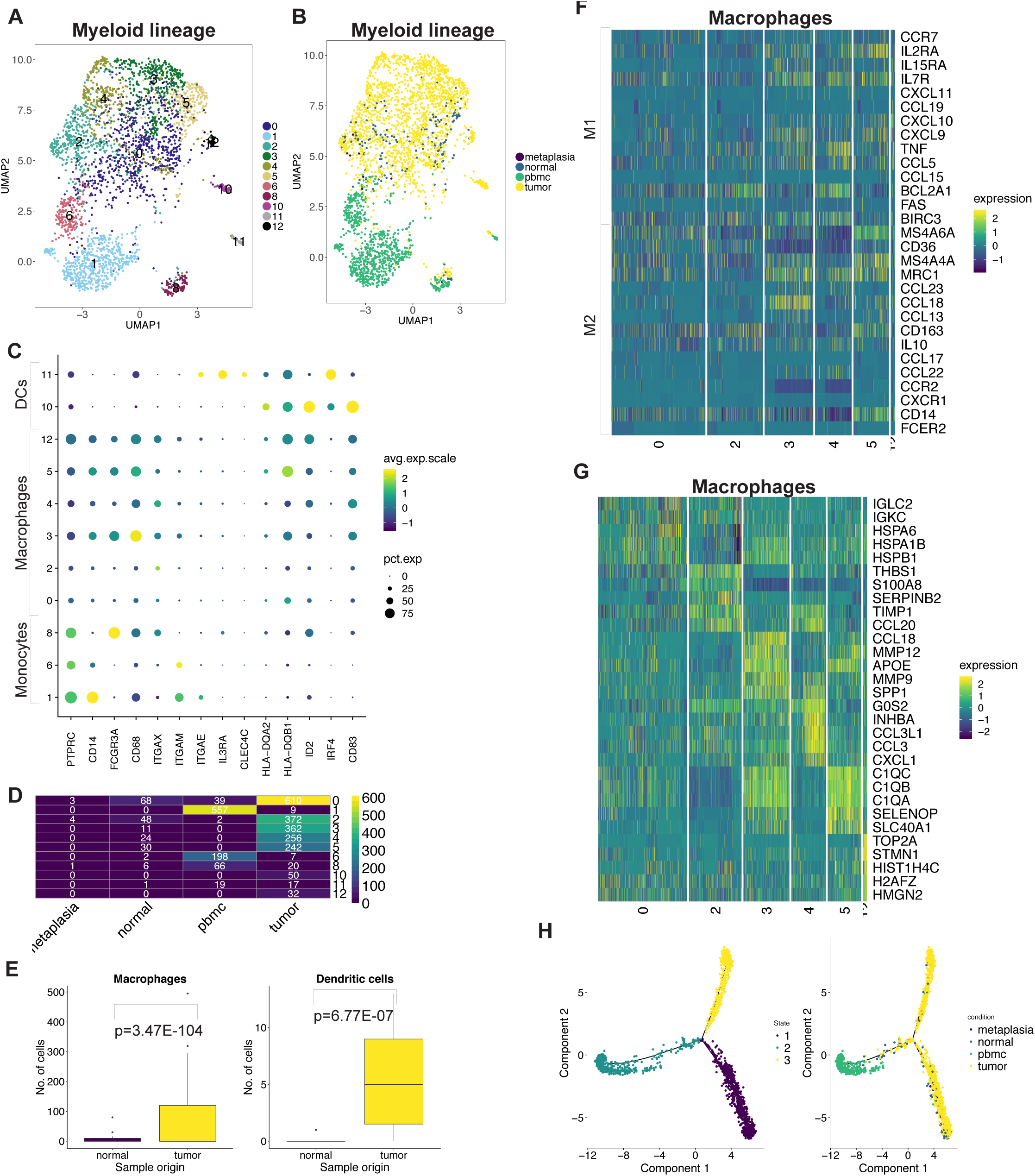
(A) UMAP representation of macrophage cells following graph-based clustering with arbitrary cluster numbers. (B) UMAP representation colored according to the sample origin. (C) Dot plot depicting expression levels of specific genes across clusters with marker-based lineage assignments. (D) Heatmap depicting number of cells identified for each cluster according the sample origin. (E) Box plots depicting proportion of macrophages from total cells derived from tumor, normal site or metaplastic with p value derived from two proportions z-test. (F) Heatmap depicting expression of M1/M2 genes from each macrophage cluster. (G) Heatmap depicting top 5 highest significantly expressed genes detected from each macrophage cluster. (H) Trajectory plots of macrophages in normal and tumor tissue with monocytes from PBMCs with cells colored by identified trajectories (left) and sample origin (right).

We examined the expression of marker genes for M1 (e.g. *CCL19*, *TNF*, *CCL5*) and M2 (e.g. *MRC1*, *CCL18*, *CCL13*, *CD163*) states across the six macrophage clusters observed in gastric tissue (15) (**Figure 3F**). Expression of M1/M2 genes did not distinguish cell types as seen in different clusters. Moreover, these genes were co-expressed in the same cluster. This result suggests that the transcriptional heterogeneity was independent of the M1/M2 classification.

We identified the differentially expressed genes (Bonferroni p < 0.05, log fold change > 0.25, genes expressed in >25% cells in a cluster) across all clusters to assess heterogeneous phenotypes (**Supplementary Table 6**). This analysis revealed that heterogeneity in the six macrophage clusters was related to significant differences (Bonferroni adjusted p < 0.05) in the expression of HSP family genes, *THBS1*, chemokines including *CCL20*, *CCL1*8, *CCL3*, matrix metalloproteinase genes, complement family genes and cell cycle regulation genes (**Figure 3G**). Clusters also showed significant differential enrichment (ANOVA, FDR p < 0.05) of hallmark gene set activity confirming their distinct transcriptional programs (**Supplementary Figure 5B**).

PBMC monocytes clustered distinctly from tumor or normal macrophages indicating transcriptional differences among the different tissue sources (**Figure 3B, D**). As it stands, conventional single cell clustering methods to do not delineate the dynamic process of cell differentiation. A new type of method called trajectory analysis, uses single cell gene expression to determine the transition among specific cell type lineages and states. Our trajectory analysis of tumor, normal tissue and PBMC macrophages yielded three different trajectory states (**Figure 3H**). Monocytes were present in a single trajectory state with few tissue macrophages. The majority of tissue macrophages differentiated along two distinct states. This results suggests that tumor infiltrating macrophages differentiate from monocytes but retain some fundamental similarities to macrophages within normal tissue.

DCs had two subclasses which demonstrated some unique transcriptional properties. One subclass had genes that define plasmacytoid DCs – these genes included *IL3RA* (CD123) and *CLEC4C* (CD303). This subclass was detected predominantly in PBMCs with only a limited number of this type of cell in the TME (**Figure 3C**, **Figure 3D**). A second DC subclass was enriched in the TME and showed significant differential expression of activated DC gene markers *CD83*, *CCR7*, *IL7R* and *ID2* (16) **(Supplementary Table S7**, **Supplementary Figure 5C**) (17). This DC subclass expressed the chemokines *CCL22*, *CCL17*, *CCL19* and *IL32* which are associated with recruitment of naïve T cells. Moreover, this subclass had differential expression of *IDO1* which is a gene marker for an immunosuppressive phenotype (18, 19). This result represented a novel gene expression program in TME infiltrating DCs not previously described.

We compared activity levels of 1,558 experimentally derived immunologic gene signatures containing the term ‘macrophage’, ‘DC’ or ‘monocyte’ **(Supplementary Methods, Supplementary Table S8**) to these gene expression profiles. The identity of monocyte and DCs were confirmed using this approach. Also, these results provided the gene set information indicating the activation phenotype of tumor specific DCs. Each macrophage cluster was enriched for gene sets derived from a variety of experimental conditions. Hence, macrophage heterogeneity and variable states likely reflect stimulus-based context with the TME.

Regulatory genes controlling gene expression of a group of genes are referred to as regulons. We identified regulons for these different transcriptional cell states using the analysis program SCENIC (5). This analysis identified transcriptional regulators such as *IRF4* in DCs and also revealed a distinct set of regulatory genes defining the various macrophage populations including *NFKB1*, *ETS2*, *CREM*, *REL*, *STAT1*, *FOXO3*, etc. **(Supplementary Table S9**). Our data provides direct *in vivo* evidence that tumor-specific macrophages exist in a continuum of stimulus-dependent functional states regulated by a specific set of genes rather than the M1/M2 paradigm. We also discovered a TME-specific gene expression program in DCs.

### TME exhausted T cells have high *CXCL13* expression and proliferation

Exhausted T cells were a prominent feature of the gastric TME compared to normal tissue. TME was also significantly enriched for Treg cells compared to normal. Initially, we identified CD4 helper T, CD8 T, NK, Treg, plasma and B cells using classic immunophenotyping markers. We filtered clusters characterized by high expression of HSP family genes and lacking lineage markers (**Supplementary Figure 6 A-D, Supplementary methods**).

Cytotoxic CD8 T lymphocytes **(CTLs)** were distributed across five different clusters indicative of transcriptionally distinct cell states. These CTL cluster states were observed among all patients (**Supplementary Figure 7A**). We examined each cluster for the dominant sample origin (normal, tumor or PBMC), expression of naïve markers (*CCR7*, *SELL*), tissue effector memory markers (*CD69*, *ITGAE*, *ITGA1*) and cytotoxic genes (*GZMB*, *GZMA*, *PRF1*, *IFNG*, *NKG7*) (**Figure 4A,B**). CTL subclasses included naïve tumor CTLs (cluster 0), effector normal CTLs (cluster 1), effector PBMC CTLs (cluster 19) and two subclasses of tumor effector CTLs (clusters 6, 22). Analysis of differential gene expression identified distinct signatures among the CTL subclasses **(Supplementary Table S10**).

**Figure 4:**
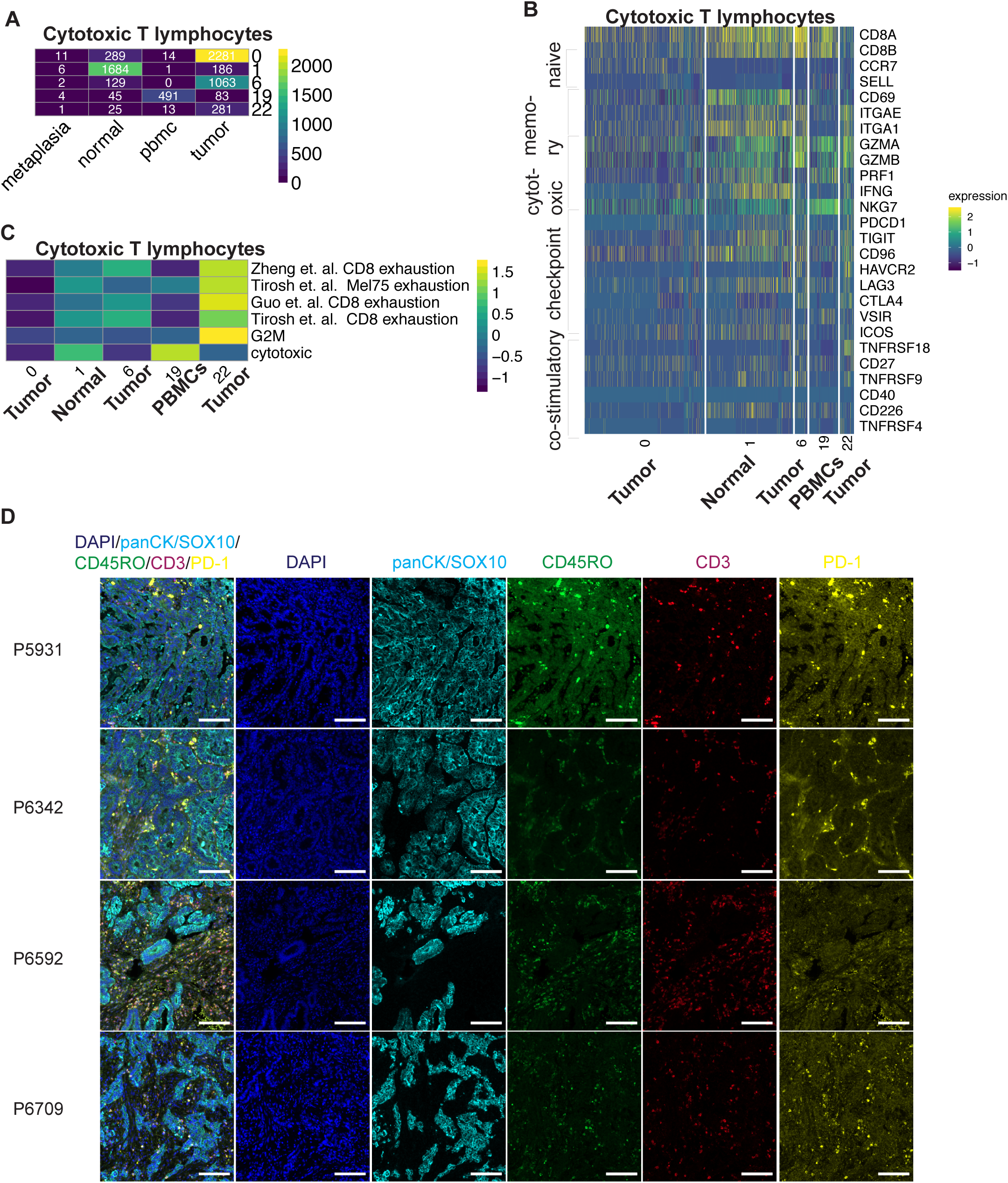
(A) Heatmap depicting number of cytotoxic T cells identified for each cluster according the sample origin. (B) Heatmap depicting expression of respective genes from each cytotoxic T cell cluster. (C) Heatmap representing average GSVA enrichment score for respective exhaustion signature for each cluster. (D) Representative images of fluorescence staining for respective markers and merged image for respective patients. Scale bar indicates 100 μm

Within cancers one observes exhausted T cells **(TEx)** which have reduced cytotoxic activity and expression of inhibitory receptors (20). In the gastric TME, both subclasses of tumor effector CTLs had low expression of *PRF1* and *IFNG* while expressing granzymes (**Figure 4B**). These genes are indicators of TEx among CTLs (21). Furthermore, these TME TEx cells expressed multiple immune checkpoints (*PDCD1*, *TIGIT*, *CD96*, *HAVCR2* (TIM3), *LAG3*, *CTLA4*, *VSIR*) indicating a degree of immunosuppression among this CTL population. These cells also expressed multiple co-stimulatory molecules (*ICOS*, *TNFRSF18* (GITR), *CD27*, *TNFRSF9*, *CD40*, *CD226*, *TNFRSF4*) (**Figure 4B**).

We compared the transcriptional profiles of the naïve tumor CTLs, effector normal CTLs, effector PBMC CTLs and two subclasses of tumor TEx cells to previously published gene signatures to further understand the tumor TEx phenotype. First, we used the gene signatures from T cell exposure to viruses. This experimental setup distinguishes the transcriptional program of naïve cells unexposed to virus, effector cells responding to the lymphocytic choriomeningitis virus **(LMCV)** and TEx cells responding to a specific clone 13 of LCMV (21).

This result confirmed our assignment of naïve, effector and TEx classes across the five clusters with greater exhaustion profiles noted in the two subclasses of the tumor specific TEx clusters (**Supplementary Figure 7B**). We additionally compared the CTL transcriptional profiles to three independently derived TEx profiles from single-cell analysis of lymphocytes in human tumors, one from mouse tumors, together with signatures for cytotoxicity and proliferation (**Figure 4C**) (22–24). This result confirmed the low-cytotoxicity, high-exhaustion phenotype in both subclasses of tumor-TEx cells.

The two subclasses of tumor-TEx significantly differed (ANOVA FDR p < 0.05) in the extent of their exhaustion and proliferation gene expression program (**Figure 4D**). The subclass with higher proliferation (cluster 22) was also associated with greater exhaustion indicative of an active immune response that has previously been associated with terminal exhaustion (25).

Immune checkpoint or costimulatory molecule gene expression was not significantly different between the two sub-classes. The second subclass (cluster 6) had lower exhaustion and proliferation plus significantly higher expression of *CXCL13* as previously identified in tumor but not in viral TEx (**Supplementary Figure 7C**) (26). Also, these cells expressed high *RBPJ*, *NR3C1* and *BATF* that are regulators of CD8 T cell fate (27). Our results thus demonstrated that effector CTLs in the TME are exhausted unlike normal tissue or PBMCs with two distinct subclasses characterized by high *CXCL13* expression or proliferation.

To verify our findings, we conducted multiplex immunofluorescence staining for pan-cytokeratin/SOX-10 (expressed in epithelial cells), CD45RO (memory T cells), CD3 (T cells) and PD-1 (exhausted T cells) in tumors from four patients where adequate tissue samples were available (**Figure 4D**, **Supplementary Figure 8A**). We detected these cell lineages in all samples including cellular sub-populations of effector T cells (CD45RO, CD3 positive) and exhausted effector T cells (PD-1, CD45RO, CD3 positive) based on co-expression analysis (**Supplementary Figure 8B**). The stromal cells and macrophages were also apparent as they lacked expression of these markers.

### Increased Tregs in the gastric TME contribute to immunosuppression

Tregs were significantly enriched in the gastric TME compared to normal gastric tissue, thus indicating an important mode of immunosuppression (**Supplementary Figure 9A, B)**. The Treg cells had two distinct subclasses distinguished by significantly higher expression of markers of proliferation (Bonferroni p < 0.05, logFC > 0.2, expressed in >25% cells) (eg. *MKI67*, *TYMS*) in one sub-class (cluster 22). In addition, Tregs expressed several immune checkpoint and costimulatory molecules representing potential targets to modulate their function (**Supplementary Figure 9C**).

CD4 T cells were represented by four subclasses. Three of these were characterized by expression of naïve markers such as the genes *CCR7*, *SELL* originating from the PBMCs or normal gastric tissue (**Supplementary Figure 9D, E**). Effector CD4 T cells (cluster 6) lacking naïve marker gene expression were found in both normal and tumor tissue (**Supplementary Figure 9D, E**). They were characterized by significantly higher expression of *GZMA*, *GZMB*, *CXCL13*, *BATF*, HLA genes (**Supplementary Figure 9E**). These cells are likely to represent follicular helper-like *CXCL13* producing CD4 cells that are associated with tertiary lymphoid structures (28, 29).

NK cells with three subclasses were detected in PBMCs (cluster 2) and both tumor and normal tissue (clusters 13, 14) (**Supplementary Figure 10A**). These cells contained a mix of rare populations of invariant NK cells, innate lymphoid cells and NK cells (**Supplementary Figure 10B**) (30). Cells in tumor and normal sites clustered together indicating transcriptional similarity. These cells in the tumor expressed cytotoxic molecules such as *GZMA*, *XCL2*, *CCL5*, *PRF1*, *CCL3*, *CCL4* indicating a potential role in mediating an anti-tumor immune response (**Supplementary Figure 10C**). Cells also expressed several inhibitory and co-stimulatory molecules including *TNFRSF18* (GITR), *CD96* and *KIR2DL4* expression representing targets for modulating their function.

B Cells from gastric TME and normal sites clustered together indicating transcriptional similarity among these cells (**Supplementary Figure 10D**). However, plasma cell clusters showed significant differences in the expression of genes encoding immunoglobulin isotypes when comparing gastric TME versus normal tissue (**Supplementary Figure 10E, F**). Specifically, plasma cells in normal tissue expressed genes encoding for IgA, those in tumor were enriched for IgG (Bonferroni p < 0.05, logFC > 0.2, expressed in >25% cells).

### Identification of jointly regulated genes of lymphocyte cell states

Our clustering analysis successfully revealed lymphocyte sub-populations driven by lineage as well as subclasses of their transcriptional cell states. Regulatory genes control group of genes with their activation or suppression, occurring as a joint unit. These regulatory genes are referred to as regulons. We identified these regulons using the analysis program SCENIC (5) **(Supplementary Table S11**). In *CXCL13*-high tumor-TEx cells we detected significant enrichment of *FOXO1* activity (ANOVA FDR p < 0.05) that is required for post antigen expansion of CD8 T cells (31). In high *CXCL13*-CD4 T cells, *BATF* activity was prominent validating their similarity to recently described *CXCL13* producing helper T cells (32). We successfully identified *FOXP3* and *BATF* gene network enrichment in Tregs confirming the accuracy of this approach. We discovered additional transcriptional regulators of Treg fate in the TME including *KDM5B*, *MAF*, *IKZF2*, *SOX4*, *BCL3*, etc. These regulons are of potential translational value given the interest in targeting epigenetics for modulating immune cell states for immunotherapy (33).

### TME reprogramming of the fibroblasts, pericytes and endothelial stroma

We discovered a transcriptional reprogramming of stromal cells in the tumor compared to normal tissue that allows the generation of a tumor-specific extracellular matrix **(ECM)**. Our analysis of the stromal cells across all samples identified fibroblasts (expressing *THY1*, *DCN*, *COL4A1*, *FAP*), endothelial cells (expressing *PECAM*, *ENG*, *VWF*, *SELE*), and pericytes (expressing *RSG5*, *PDGFRB*) (**Figure 5A, 5C**). The observed heterogeneity among fibroblasts is likely to be driven by patient specific factors (**Supplementary Figure 11A**). Endothelial clusters additionally differed in the expression of genes encoding secretory factors (e.g. *ESM1*, *ANGPT2*), tip cell markers (e.g. *COL4A1*, *COL4A2*, *DLL4*, *MARCKSL1*) and stalk cell markers (*ACKR1*, *CD36*, *SELP*, *VWF*) (**Supplementary Figure 11B, 11C)** (34).

**Figure 5:**
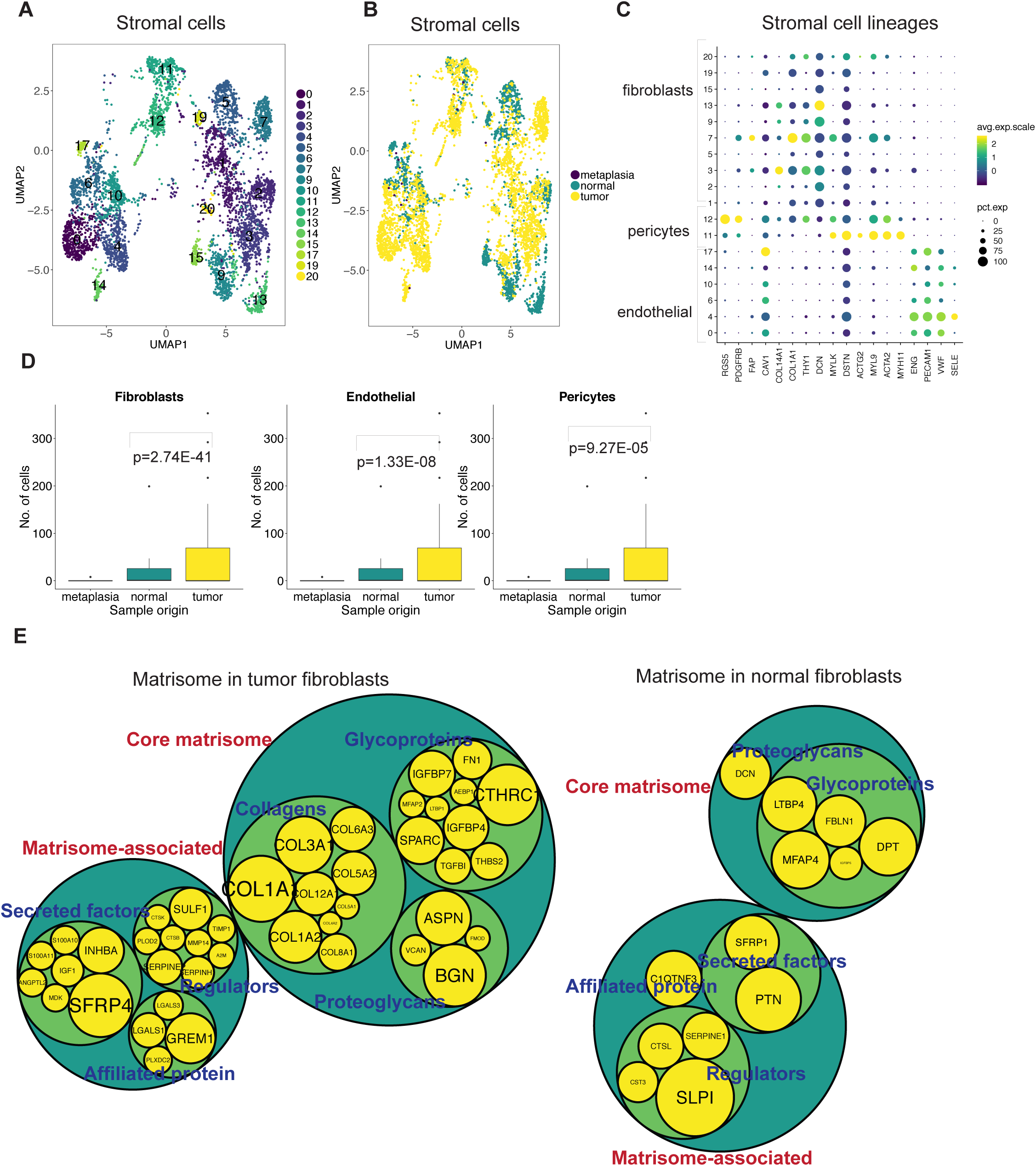
(A) UMAP representation of stromal cells following graph-based clustering with arbitrary cluster numbers and (B) colored according to the sample origin. (C) Dot plot depicting expression levels of specific genes across clusters with marker-based lineage assignments. (D) Box plots depicting proportion of fibroblasts, pericytes or endothelial cells from total cells derived from tumor, normal or metaplastic site with p value derived from two proportions z-test. (E) Comparison of differentially expressed genes in tumor or normal fibroblasts to the genes of the matrisome program. Size of gene level circles is proportional to the logFC.

Importantly, all three cell types were enriched in tumor tissue compared to normal (**Figure 5B,5D**). Stromal cells are responsible for the production and maintenance of ECM that provides mechanical support to cells and also influences their growth by interactions. Hence, these differences in number can impact the characteristics of ECM in tumor compared to normal tissue.

Genes encoding for components or regulators of the ECM have previously been identified as the ‘matrisome’ (35, 36). It consists of core factors such as collagens, proteoglycans and ECM glycoproteins that make up the ECM as well as an associated program that consists of ECM regulators, secretory factors and ECM-affiliated proteins. To understand the phenotypical differences in stromal cells found in normal or tumor tissue, we compared their significantly differentially expressed genes (Bonferroni adjusted p < 0.05, logFC > 0.25 and expressed in > 25% cells) to the matrisome gene expression program. Tumor-specific fibroblasts, pericytes and endothelial cells expressed diverse ECM components, including glycoproteins, collagens and proteoglycans, as well as ECM regulators, affiliated proteins and secretory factors as compared to normal stromal cells (**Figure 5E**, **Supplementary Table S12**). Additionally, fibroblasts in tumors had significant overexpression of *ACTA2* compared to normal tissue, indicative of their contractile ability (**Supplementary Figure 11D**).

Stromal cells at tumor or normal sites had significantly different regulatory genes or regulons (ANOVA FDR < 0.05) **(Supplementary Table S13**). For example, tumor-specific endothelial cells had greater activity of *SOX18* and *SOX7* which are known regulators of a variety of endothelial cell processes (37). Tumor-specific fibroblasts had high activity of *EGR2* that can influence fibrosis (38). Tumor-specific pericytes were enriched for *FOXF2* activity that is known to regulate pericyte differentiation (39). Hence, our approach distinguished differences in both the gene expression program and its regulators between tumor and normal stromal cells.

### TME specific cellular communication has the potential to influence cell states

We discovered a TME-specific intercellular communications network that can potentially affect cellular behavior. First, we identified receptor-ligand interactions between different cell types using CellPhoneDB, which infers statistically significant interactions from a comprehensive data catalogue. We then compared these networks between tumor and normal tissue to generate an interactome specific to the TME, which was not present at significant levels in normal tissue (**Figure 6A**, **Supplementary Table 14**).

**Figure 6:**
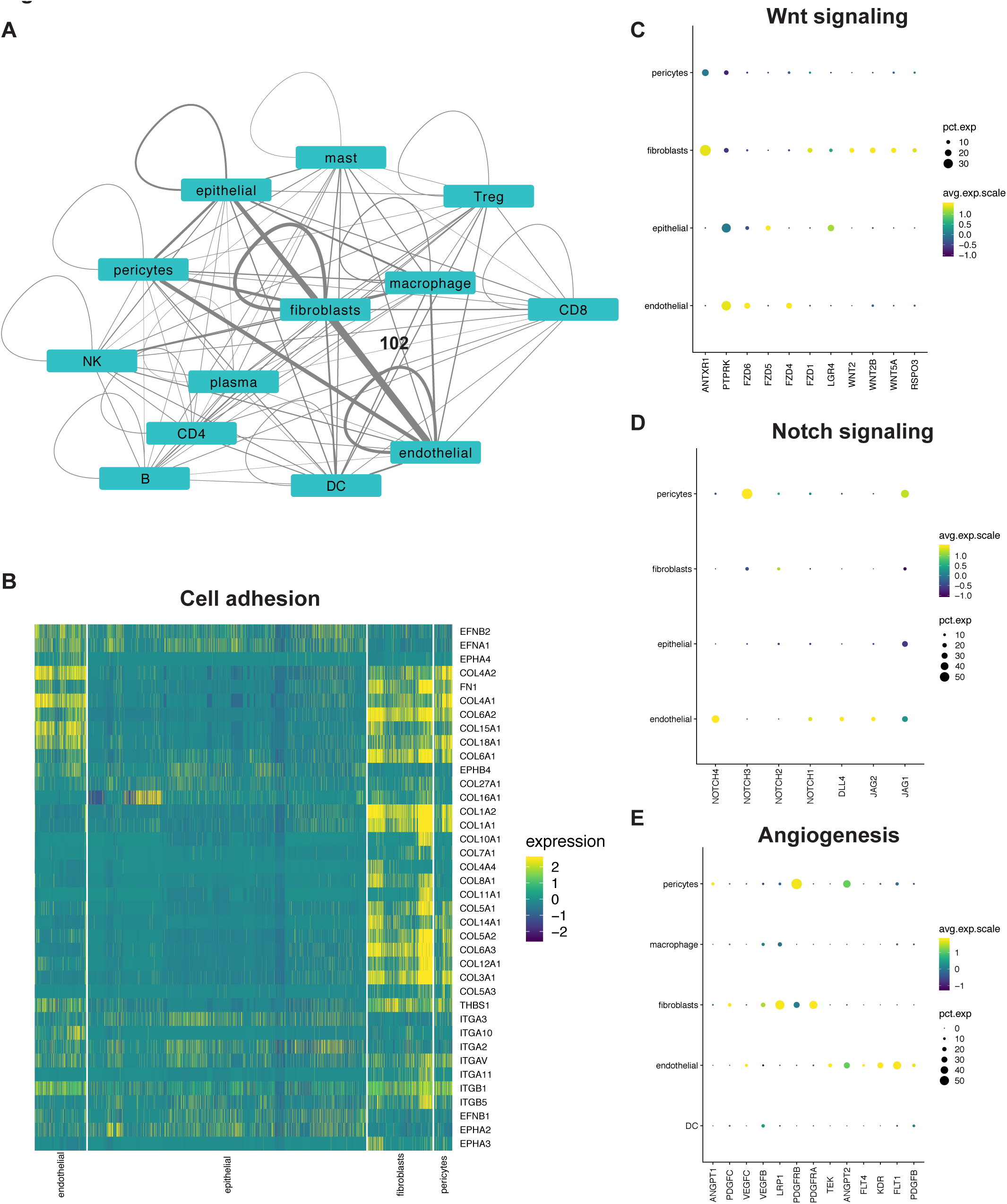
(A) Network depicting interactions between various cell types in tumors. Each node is a cell type and scaled edges represent the number of statistically significant detected interactions. Scale: fibroblast and endothelial edge = 102 interactions (B) Heatmap and (C-E) dot plots depicting the expression of respective genes in specific cell types.

Stromal cells were among the most prolific interactors. Prominent communication between epithelial cells, fibroblasts, pericytes and endothelial cells occurred through various integrin receptor interactions with collagen, fibronectin and *THBS1* ligands (**Figure 6B**). We also detected bidirectional interactions between ephrin receptor family and ephrin ligands in epithelial and endothelial cells that can influence cell phenotypes (40). Among growth factor signaling that can promote proliferation and survival of cancer cells, we could detect expression of *EGFR* and *MET* receptors on epithelial cells together with respective ligand expression on stromal cells. We also detected significant *EGFR* signaling interactions in fibroblasts.

Fibroblasts were a prominent source of Wnt ligands with expression of corresponding receptors on tumor epithelial cells, endothelial cells, fibroblasts and pericytes. This included a *LGR4* – *RSPO3* interaction that has the potential to regulate stemness (41) (**Figure 6C**), validating our previous discovery of fibroblast derived *RSPO3* in an organoid model of gastric cancer (42).

Autocrine Notch signaling was evident in endothelial cells, a known regulator of angiogenesis (43), together with paracrine support from tumor epithelium and fibroblasts. Interactions promoting Notch signaling, which can regulate their transition to myofibroblasts (44), were also significant in fibroblasts (**Figure 6D**). Angiogenic receptors *KDR*, *FLT1*, *FLT4*, *PDGFB*, *TEK* on endothelial cells and pericytes had significant autocrine and paracrine interactions with their respective ligands (**Figure 6E**). Among the interactome were 19 cytokines including chemokines, interleukins, tumor necrosis factors (TNFs) and their corresponding receptors that can influence immune cell fates.

## DISCUSSION

We leveraged paired distal normal tissue and PBMCs in our study to analyze the cellular dysregulation and biological changes in the GC TME. With single cell gene expression analysis, we demonstrated that GC TME leads to a series of dramatic cellular changes compared to matched normal stomach mucosa. Specifically, we noted increases in cell numbers of fibroblasts, endothelial cells, pericytes and Tregs in the TME. We also identified transcriptional cell states unique to the TME. This included two subclasses of exhausted CTLs unique to the TME characterized by *CXCL13* expression or increased proliferation and TME-specific DCs. We also generated gene expression profiles for tumor associated macrophages that consisted of six heterogenous subclasses not confined to a binary M1/M2 classification. We demonstrated that stromal cells in the TME have a gene expression program not found in normal tissue that encodes for a specific extracellular matrix composition. These results identified novel gene regulatory networks and intercellular communication networks across these TME specific cell types and states.

We validated previously described changes in normal, metaplastic and tumor epithelial cells (12, 45) and were additionally able to elucidate intra-tumor heterogeneity by examining the activity of various cancer promoting mechanisms within the tumor cells. The diversity in activation profiles suggests that targeting multiple clones with different combination strategies may be necessary to eradicate them completely.

Immunosuppression in GC TME was evident by the increased proportion of Tregs compared to normal tissue. We identified several checkpoint and costimulatory molecules on these cells, and their transcriptional regulators. These regulators can be investigated to understand Treg biology and to derive therapeutic targets. Indeed, recent evidence indicates that anti-CTLA4 activity is a consequence of Treg depletion in the TME (46). We detected expression of multiple immune checkpoints on cytotoxic T cells similar to other studies (22, 47). These checkpoints were also detected on helper T and Treg subsets. Thus, it is important to understand the effects of immune checkpoint blockade on distinct T cell subpopulations that express the same target. Our analysis reveals molecules and transcriptional regulators responsible for these states. Plasma cells in tumor tissue expressed IgGs rather than IgAs that were detected in paired normal tissue. IgGs have been associated with a pro-cancer role by influencing myeloid cell Fc-receptors (48).

We demonstrated that tumor immune cells are transcriptionally closer to normal tissue counterparts than PBMCs. Our trajectory analysis shows that the majority of tissue macrophages are distinct from monocytes, supporting their different origins (49). Application of reference profiles derived from PBMCs to tumor infiltrating immune cells should also be interpreted with caution.

Interactome analysis demonstrated pro-tumor effects of TME components and also the influence of cancer cells on the TME. Tumor-specific interactome represents potential treatment targets to inhibit cancer proliferation, overcome the immunosuppressive microenvironment and restore the cancer immunity cycle. Additionally, while some targets such as Wnt inhibition have previously been regarded only in the context of tumor epithelial cells, our analysis demonstrates that this might have implications for the TME.

Our study did not consider spatial context and might be affected by the dissociation process. For immune cells in particular, the use of dual single-cell proteomics and transcriptomics is likely to provide a more refined analysis of immune cell sub-types (50).

## DISCLOSURE OF POTENTIAL CONFLICTS OF INTEREST

None to disclose.

## AUTHORS’ CONTRIBUTIONS

AS was involved in conception and design of the study, development of methodology, acquisition of data, analysis and interpretation of data and writing of the manuscript. SG was involved in analysis and interpretation of data. BTL and JC were involved in the development of methodology and acquisition of data. CS was involved in the interpretation of data. RH and GP were involved in design of the study and material support. HPJ oversaw the conception and design of the study, interpretation of data and writing of the manuscript.

## ACKNOWLEDGEMENTS

We are grateful to all patients who participated in the study. We thank Christine Handy, Alison Almeda, Christina Wood-Bouwens for assistance in sample collection and documentation and Ann Renschler for assistance in sequencing. We thank Dr. Katir Patel from Ultivue, Inc. for assistance in multiplex immunofluorescence staining. This work was supported by US National Institutes of Health grants P01HG000205 (HPJ, CWB and SMG), R01HG006137 (HPJ), U01CA217875 (HPJ and AS). HPJ also received support from the American Cancer Society (124571-RSG-13-297-01), the Clayville Foundation and the Gastric Cancer Foundation.

## REFERENCES

1. Bray F, Ferlay J, Soerjomataram I, Siegel RL, Torre LA, Jemal A. Global cancer statistics 2018: GLOBOCAN estimates of incidence and mortality worldwide for 36 cancers in 185 countries. CA Cancer J Clin 2018;68(6):394–424 doi 10.3322/caac.21492.

2. Cislo M, Filip AA, Arnold Offerhaus GJ, Cisel B, Rawicz-Pruszynski K, Skierucha M, et al. Distinct molecular subtypes of gastric cancer: from Lauren to molecular pathology. Oncotarget 2018;9(27):19427–42 doi 10.18632/oncotarget.24827.

3. Egeblad M, Nakasone ES, Werb Z. Tumors as organs: complex tissues that interface with the entire organism. Dev Cell 2010;18(6):884–901 doi 10.1016/j.devcel.2010.05.012.

4. Shitara K, Ozguroglu M, Bang YJ, Di Bartolomeo M, Mandala M, Ryu MH, et al. Pembrolizumab versus paclitaxel for previously treated, advanced gastric or gastro-oesophageal junction cancer (KEYNOTE-061): a randomised, open-label, controlled, phase 3 trial. Lancet 2018;392(10142):123–33 doi 10.1016/S0140-6736(18)31257-1.

5. Aibar S, Gonzalez-Blas CB, Moerman T, Huynh-Thu VA, Imrichova H, Hulselmans G, et al. SCENIC: single-cell regulatory network inference and clustering. Nat Methods 2017;14(11):1083–6 doi 10.1038/nmeth.4463.

6. Qiu X, Mao Q, Tang Y, Wang L, Chawla R, Pliner HA, et al. Reversed graph embedding resolves complex single-cell trajectories. Nat Methods 2017;14(10):979–82 doi 10.1038/nmeth.4402.

7. Vento-Tormo R, Efremova M, Botting RA, Turco MY, Vento-Tormo M, Meyer KB, et al. Single-cell reconstruction of the early maternal-fetal interface in humans. Nature 2018;563(7731):347–53 doi 10.1038/s41586-018-0698-6.

8. Butler A, Hoffman P, Smibert P, Papalexi E, Satija R. Integrating single-cell transcriptomic data across different conditions, technologies, and species. Nat Biotechnol 2018;36(5):411–20 doi 10.1038/nbt.4096.

9. McInnes L, Healy J. UMAP: uniform manifold approximation and projection for dimension reduction. ArXiv2018.

10. Alquicira-Hernández J, Sathe A, Ji H, Nguyen Q, Powell JE. scPred: Cell type prediction at single-cell resolution. bioRxiv2018.

11. Cancer Genome Atlas Research N. Comprehensive molecular characterization of gastric adenocarcinoma. Nature 2014;513(7517):202–9 doi 10.1038/nature13480.

12. Zhang P, Yang M, Zhang Y, Xiao S, Lai X, Tan A, et al. Dissecting the Single-Cell Transcriptome Network Underlying Gastric Premalignant Lesions and Early Gastric Cancer. Cell Rep 2019;27(6):1934–47 e5 doi 10.1016/j.celrep.2019.04.052.

13. Muller S, Cho A, Liu SJ, Lim DA, Diaz A. CONICS integrates scRNA-seq with DNA sequencing to map gene expression to tumor sub-clones. Bioinformatics 2018;34(18):3217–9 doi 10.1093/bioinformatics/bty316.

14. Schumacher SE, Shim BY, Corso G, Ryu MH, Kang YK, Roviello F, et al. Somatic copy number alterations in gastric adenocarcinomas among Asian and Western patients. PLoS One 2017;12(4):e0176045 doi 10.1371/journal.pone.0176045.

15. Martinez FO, Gordon S, Locati M, Mantovani A. Transcriptional profiling of the human monocyte-to-macrophage differentiation and polarization: new molecules and patterns of gene expression. J Immunol 2006;177(10):7303–11 doi 10.4049/jimmunol.177.10.7303.

16. Collin M, McGovern N, Haniffa M. Human dendritic cell subsets. Immunology 2013;140(1):22–30 doi 10.1111/imm.12117.

17. Villani AC, Satija R, Reynolds G, Sarkizova S, Shekhar K, Fletcher J, et al. Single-cell RNA-seq reveals new types of human blood dendritic cells, monocytes, and progenitors. Science 2017;356(6335) doi 10.1126/science.aah4573.

18. Pietila TE, Veckman V, Lehtonen A, Lin R, Hiscott J, Julkunen I. Multiple NF-kappaB and IFN regulatory factor family transcription factors regulate CCL19 gene expression in human monocyte-derived dendritic cells. J Immunol 2007;178(1):253–61 doi 10.4049/jimmunol.178.1.253.

19. Mellor AL, Munn DH. IDO expression by dendritic cells: tolerance and tryptophan catabolism. Nat Rev Immunol 2004;4(10):762–74 doi 10.1038/nri1457.

20. Wherry EJ, Kurachi M. Molecular and cellular insights into T cell exhaustion. Nat Rev Immunol 2015;15(8):486–99 doi 10.1038/nri3862.

21. Wherry EJ, Ha SJ, Kaech SM, Haining WN, Sarkar S, Kalia V, et al. Molecular signature of CD8+ T cell exhaustion during chronic viral infection. Immunity 2007;27(4):670–84 doi 10.1016/j.immuni.2007.09.006.

22. Tirosh I, Izar B, Prakadan SM, Wadsworth MH, 2nd, Treacy D, Trombetta JJ, et al. Dissecting the multicellular ecosystem of metastatic melanoma by single-cell RNA-seq. Science 2016;352(6282):189–96 doi 10.1126/science.aad0501.

23. Guo X, Zhang Y, Zheng L, Zheng C, Song J, Zhang Q, et al. Global characterization of T cells in non-small-cell lung cancer by single-cell sequencing. Nat Med 2018;24(7):978–85 doi 10.1038/s41591-018-0045-3.

24. Zheng C, Zheng L, Yoo JK, Guo H, Zhang Y, Guo X, et al. Landscape of Infiltrating T Cells in Liver Cancer Revealed by Single-Cell Sequencing. Cell 2017;169(7):1342–56 e16 doi 10.1016/j.cell.2017.05.035.

25. Miller BC, Sen DR, Al Abosy R, Bi K, Virkud YV, LaFleur MW, et al. Subsets of exhausted CD8(+) T cells differentially mediate tumor control and respond to checkpoint blockade. Nat Immunol 2019;20(3):326–36 doi 10.1038/s41590-019-0312-6.

26. Thommen DS, Schumacher TN. T Cell Dysfunction in Cancer. Cancer Cell 2018;33(4):547–62 doi 10.1016/j.ccell.2018.03.012.

27. Yu B, Zhang K, Milner JJ, Toma C, Chen R, Scott-Browne JP, et al. Epigenetic landscapes reveal transcription factors that regulate CD8(+) T cell differentiation. Nat Immunol 2017;18(5):573–82 doi 10.1038/ni.3706.

28. Gu-Trantien C, Loi S, Garaud S, Equeter C, Libin M, de Wind A, et al. CD4(+) follicular helper T cell infiltration predicts breast cancer survival. J Clin Invest 2013;123(7):2873–92 doi 10.1172/JCI67428.

29. Kobayashi S, Watanabe T, Suzuki R, Furu M, Ito H, Ito J, et al. TGF-beta induces the differentiation of human CXCL13-producing CD4(+) T cells. Eur J Immunol 2016;46(2):360–71 doi 10.1002/eji.201546043.

30. Bezman NA, Kim CC, Sun JC, Min-Oo G, Hendricks DW, Kamimura Y, et al. Molecular definition of the identity and activation of natural killer cells. Nat Immunol 2012;13(10):1000–9 doi 10.1038/ni.2395.

31. Hedrick SM, Hess Michelini R, Doedens AL, Goldrath AW, Stone EL. FOXO transcription factors throughout T cell biology. Nat Rev Immunol 2012;12(9):649–61 doi 10.1038/nri3278.

32. Yoshitomi H, Kobayashi S, Miyagawa-Hayashino A, Okahata A, Doi K, Nishitani K, et al. Human Sox4 facilitates the development of CXCL13-producing helper T cells in inflammatory environments. Nat Commun 2018;9(1):3762 doi 10.1038/s41467-018-06187-0.

33. Dunn J, Rao S. Epigenetics and immunotherapy: The current state of play. Mol Immunol 2017;87:227–39 doi 10.1016/j.molimm.2017.04.012.

34. Zhao Q, Eichten A, Parveen A, Adler C, Huang Y, Wang W, et al. Single-Cell Transcriptome Analyses Reveal Endothelial Cell Heterogeneity in Tumors and Changes following Antiangiogenic Treatment. Cancer Res 2018;78(9):2370–82 doi 10.1158/0008-5472.CAN-17-2728.

35. Naba A, Clauser KR, Hoersch S, Liu H, Carr SA, Hynes RO. The matrisome: in silico definition and in vivo characterization by proteomics of normal and tumor extracellular matrices. Mol Cell Proteomics 2012;11(4):M111 014647 doi 10.1074/mcp.M111.014647.

36. Hynes RO, Naba A. Overview of the matrisome--an inventory of extracellular matrix constituents and functions. Cold Spring Harb Perspect Biol 2012;4(1):a004903 doi 10.1101/cshperspect.a004903.

37. De Val S, Black BL. Transcriptional control of endothelial cell development. Dev Cell 2009;16(2):180–95 doi 10.1016/j.devcel.2009.01.014.

38. Bhattacharyya S, Wu M, Fang F, Tourtellotte W, Feghali-Bostwick C, Varga J. Early growth response transcription factors: key mediators of fibrosis and novel targets for anti-fibrotic therapy. Matrix Biol 2011;30(4):235–42 doi 10.1016/j.matbio.2011.03.005.

39. Reyahi A, Nik AM, Ghiami M, Gritli-Linde A, Ponten F, Johansson BR, et al. Foxf2 Is Required for Brain Pericyte Differentiation and Development and Maintenance of the Blood-Brain Barrier. Dev Cell 2015;34(1):19–32 doi 10.1016/j.devcel.2015.05.008.

40. Lisabeth EM, Falivelli G, Pasquale EB. Eph receptor signaling and ephrins. Cold Spring Harb Perspect Biol 2013;5(9) doi 10.1101/cshperspect.a009159.

41. Barker N, Tan S, Clevers H. Lgr proteins in epithelial stem cell biology. Development 2013;140(12):2484–94 doi 10.1242/dev.083113.

42. Chen J, Lau BT, Andor N, Grimes SM, Handy C, Wood-Bouwens C, et al. Single-cell transcriptome analysis identifies distinct cell types and niche signaling in a primary gastric organoid model. Sci Rep 2019;9(1):4536 doi 10.1038/s41598-019-40809-x.

43. Siekmann AF, Lawson ND. Notch signalling and the regulation of angiogenesis. Cell Adh Migr 2007;1(2):104–6.

44. Dees C, Tomcik M, Zerr P, Akhmetshina A, Horn A, Palumbo K, et al. Notch signalling regulates fibroblast activation and collagen release in systemic sclerosis. Ann Rheum Dis 2011;70(7):1304–10 doi 10.1136/ard.2010.134742.

45. Companioni O, Sanz-Anquela JM, Pardo ML, Puigdecanet E, Nonell L, Garcia N, et al. Gene expression study and pathway analysis of histological subtypes of intestinal metaplasia that progress to gastric cancer. PLoS One 2017;12(4):e0176043 doi 10.1371/journal.pone.0176043.

46. Tang F, Du X, Liu M, Zheng P, Liu Y. Anti-CTLA-4 antibodies in cancer immunotherapy: selective depletion of intratumoral regulatory T cells or checkpoint blockade? Cell Biosci 2018;8:30 doi 10.1186/s13578-018-0229-z.

47. Andor N, Simonds EF, Czerwinski DK, Chen J, Grimes SM, Wood-Bouwens C, et al. Single-cell RNA-Seq of lymphoma cancers reveals malignant B cell types and co-expression of T cell immune checkpoints. Blood 2018 doi 10.1182/blood-2018-08-862292.

48. Garaud S, Zayakin P, Buisseret L, Rulle U, Silina K, de Wind A, et al. Antigen Specificity and Clinical Significance of IgG and IgA Autoantibodies Produced in situ by Tumor-Infiltrating B Cells in Breast Cancer. Front Immunol 2018;9:2660 doi 10.3389/fimmu.2018.02660.

49. Davies LC, Taylor PR. Tissue-resident macrophages: then and now. Immunology 2015;144(4):541–8 doi 10.1111/imm.12451.

50. Stoeckius M, Hafemeister C, Stephenson W, Houck-Loomis B, Chattopadhyay PK, Swerdlow H, et al. Simultaneous epitope and transcriptome measurement in single cells. Nat Methods 2017;14(9):865–8 doi 10.1038/nmeth.4380.

