## Supplementary information for "Single cell genomic characterization reveals the cellular reprogramming of the gastric tumor microenvironment"

**Title:**

**Institutions:**

[1] Division of Oncology, Department of Medicine, Stanford University School of Medicine, Stanford, CA, United States

[2] Stanford Genome Technology Center, Stanford University, Palo Alto, CA, United States

[3] Department of Surgery, Stanford University, Stanford, CA 94305, United States

[4] Division of Gastroenterology and Hepatology, Department of Medicine, Stanford University School of Medicine, Stanford, CA 94305, USA

[5] Department of Pathology, Stanford University School of Medicine, Stanford, CA USA

**Corresponding author:**

Hanlee P. Ji

#### SUPPLEMENTARY METHODS

**Histopathology:** For P5931, P6207, P6342, P6709, P6592 tissue was additionally fixed in 10% formalin for 24 hours at room temperature. Paraffin embedding and processing with hematoxylin and eosin staining was conducted by the Human Pathology Histology Services core facility at Stanford University. Slides were reviewed by a board-certified pathologist. We additionally reviewed clinical histopathology reports for all patients. MSI/MSS status was determined from these reports where immunohistological staining was performed on paraffin embedded tissue sections using standard protocols for MLH1, MSH2, MSH6 and PMS2.

**Multiplex immunofluorescence staining and analysis:** FFPE slides were deparaffinized and rehydrated by sequential five-minute washes in Histoclear twice, 100% ethanol twice, 90% ethanol, 70% ethanol, 50% ethanol, 30% ethanol followed by distilled water. Antigen retrieval was performed in an electronic pressure cooker for 20 minutes in a Tris-EDTA buffer with pH 9.0. Multiplex immunofluorescence staining was performed using the UltiMapper I/O PD-1 kit (Ultivue, Inc.) as per manufacturer's protocol. Whole slide images were obtained by scanning slides using Zeiss Axio Scan.Z1 (Carl Zeiss AG, Germany) with a 20x objective and ZEN software (Zeiss). Image analysis was conducted using QuPath (v0.1.2) on five different regions of interest per image (1). Cells were detected using default parameters by using the DAPI channel and changing size limits to 5 to 400 μm. Number of cells in each ROI was 2968, 2276, 2575, 3381, 1427 for P5931, 3930, 2883, 3323, 2682, 3094 for P6342, 1953, 2081, 2288, 3368, 2589 for P6592, 5100, 4940, 5880, 6798, 5465 for P6709. A classifier was created in QuPath to identify negative or positive cells for each single stain or double, triple or quadruple stains. Thresholds for channel positivity were set for each channel by visual inspection of staining and distribution of staining intensity using histograms. Process was iterated across all regions of interest and all images using the same thresholds for each individual channel. Resulting staining intensity distributions per cell and cell identities were processed in R and visualized as violin plots.

**Computational identification of doublets:** For Seurat data objects containing greater than 1000 cells, we identified doublets using DoubletFinder (v2.0.1) (2). The 'paramSweep' and 'summarizeSweep' functions were used with default settings to determine the 'pK' value - this defines the PC neighborhood size of merged real-artificial data. The 'pN' value was set to 0.25 as the proportion of artificial doublets. The proportion of homotypic doublets were identified using the 'modelHomotypic' function using marker-based lineage annotations for each cluster as described in 'Methods' section. Singlet versus doublet classifications were obtained using 'doubletFinder' function with two settings. First, 'nExP' was set to expected doublet rate according to Chromium Single Cell 3' v2 reagents kits user guide (10x Genomics). Second, this rate was adjusted to the proportion of homotypic doublets. Cells classified as a doublet with either setting were assigned a doublet designation.

**Regulome analysis:** To understand gene regulatory networks in each cell we used SCENIC (v1.1.1.5) (3), together with dependencies AUCell (v1.5.5), Rcistarget (v1.3.4) and Genie3 (v1.5.4) respectively. Expression matrix from the data slot of the Seurat object of interest was used as input to the 'initializeScenic' function together with cistarget databases

'hg38_refseqr80_500bp_up_and_100bp_down_tss.mc9nr.feather',

hg38_regseq-r80_10kb_up_and_down_tss.mc9nr.feather'

downloaded from https://resources.aertslab.org/cistarget/. Expression matrix was filtered for genes expressed with greater than 3 UMIs in 1% of cells and genes detected in at least 1% of cells. Positive and negative associations were identified by determining Spearman correlation on expression matrix. Regulators were identified using ‘runGenie3’ function with default parameters. Gene regulatory networks were scored using default parameters in ‘runSCENIC_1_coexNetwork2modules’, ‘runSCENIC_2_createRegulons’ and ‘runSCENIC_3_scoreCells’. Regulon activity was determined for each cluster by averaging the AUC values per cell. Regulon activities that were significantly different across clusters after applying ANOVA with an FDR corrected $p < 0.05$ were represented as heatmaps or tables scaled by rows.

**Pathway or gene set quantification using gene set variation analysis (GSVA):** C2 Reactome, C6 oncogenic signatures , C7 immunologic signatures gene sets were downloaded from MSigDB v 6.2 at http://software.broadinstitute.org/gsea/msigdb/genesets.jsp and read using ‘getGmt’ function in GSEAbase (v1.40.1) (4). T cell exhaustion and cytotoxic signatures were obtained from literature (5-7). Hallmark G2M checkpoint gene set was used to assess proliferation. Expression matrix derived from the data slot of the Seurat object was compared to gene sets by calculating enrichment scores for each cell using GSVA (v1.26.0) with function ‘gsva’ and parameters ‘kcdf’ and ‘mx.diff’ set to Gaussian and true respectively. Average enrichment score was determined for each cluster. Significantly different scores across clusters were determined by ANOVA with an FDR corrected p value < 0.05 and represented as heatmaps or tables scaled by rows. Default clustering parameters in ‘pheatmap’ function were used to cluster columns. For GSVA analysis of intra-tumoral heterogeneity in epithelial cells, we analyzed clusters containing greater than ten cells.

**Copy number analysis:** CONICSmat (v0.0.0.1) (8) was used to infer copy number changes at chromosome arm level based on direction of changes in gene expression. Default parameters were used for all functions unless otherwise indicated. Information of gene positions was added to the Seurat data expression matrix using ‘getGenePositions’ followed by filtering with ‘filterMatrix’ and normalization with ‘calcNormFactors’. Mixture models for z-score of centered gene expression across all cells were calculated using ‘plotAll’. For each cell, posterior probabilities were binarized as <0.9 or >0.9 for absence or presence of alterations respectively and represented as a heatmap or table.

**Trajectory analysis:** Single-cell gene expression trajectories were constructed using Moncole (v2.6.4) by importing Seurat data using the ‘importCDS’ function. Size factors and dispersion values were calculated using ‘estimateSizeFactors’ and ‘estimateDispersions’ functions respectively and filtered for genes detected using a threshold of 0.1 mean expression. These genes were used as input to ‘setorderingfilter’ function, dimensionality reduction was conducted using the ‘reduceDimension’ function with the DDRTree method and cells were placed onto a pseudotime trajectory using ‘orderCells’.

**Prediction of cell types as tumor or normal epithelium:** We constructed a training dataset composed half of normal and tumor epithelial cells selected using a seed of 51. We then used scPred (v0.0.0.9000) (9) to conduct PCA analysis on the first 10 principal components using the ‘eigenDecompose’ function and determined the most informative components using the ‘getFeatureSpace’ function. Model was built on training dataset using the ‘trainModel’ function.

Conditional class probabilities for the test data to belong to the tumor or normal class were obtained using ‘scPredict’ with a threshold of 0.55 and ‘getPredictions’.

**‘Gating’ of cells in lymphocyte clusters 6 and 22:** Cluster 22 cells that had normalized gene expression of *MS4A1* (CD20) greater than 0.001 were first designated as B cells. From the remaining, cells with normalized gene expression of *CD4* or *FOXP3* greater than 0.001 were regarded as Treg cells. All other cells were designated as CD8 cells. Cluster 6 cells with normalized gene expression of *CD8A* or *CD8B* greater than 0.001 were regarded as CD8 cells and all other cells were designated at CD4 T cells.

**Interactions between cell types:** We extracted the expression matrix of each cell lineage for either cells derived from gastric tumor or gastric normal tissue. Expression matrix together with cell type metadata was used as input to cellphonedb (release 0.0.6) (10) run with 'statistical analysis method' in Python (v2.7.13). This identified statistically significant interactions ($p < 0.05$) between receptors and ligands against a null distribution using a curated receptor-ligand database.

**Additional analysis:** Additional analysis or visualization was conducted using R packages tidyverse (v1.2.1), gplots (v3.0.1.1), broom (v0.5.2) and viridis (0.5.1) in R v3.4.4.

#### REFERENCES


1. Bankhead P, Loughrey MB, Fernandez JA, Dombrowski Y, McArt DG, Dunne PD*, et al.* QuPath: Open source software for digital pathology image analysis. Sci Rep **2017**;7(1):16878 doi 10.1038/s41598-017-17204-5.
2. McGinnis CS, Murrow LM, Gartner ZJ. DoubletFinder: Doublet Detection in Single-Cell RNA Sequencing Data Using Artificial Nearest Neighbors. Cell Syst **2019**;8(4):329-37 e4 doi 10.1016/j.cels.2019.03.003.
3. Aibar S, Gonzalez-Blas CB, Moerman T, Huynh-Thu VA, Imrichova H, Hulselmans G*, et al.* SCENIC: single-cell regulatory network inference and clustering. Nat Methods **2017**;14(11):1083-6 doi 10.1038/nmeth.4463.
4. Morgan M, Falcon S, Gentleman R. GSEABase: Gene set enrichment data structures and methods. R package version 1.46.0. 2019.
5. Tirosh I, Izar B, Prakadan SM, Wadsworth MH, 2nd, Treacy D, Trombetta JJ*, et al.* Dissecting the multicellular ecosystem of metastatic melanoma by single-cell RNA-seq. Science **2016**;352(6282):189-96 doi 10.1126/science.aad0501.
6. Guo X, Zhang Y, Zheng L, Zheng C, Song J, Zhang Q*, et al.* Global characterization of T cells in non-small-cell lung cancer by single-cell sequencing. Nat Med **2018**;24(7):978-85 doi 10.1038/s41591-018-0045-3.
7. Zheng C, Zheng L, Yoo JK, Guo H, Zhang Y, Guo X*, et al.* Landscape of Infiltrating T Cells in Liver Cancer Revealed by Single-Cell Sequencing. Cell **2017**;169(7):1342-56 e16 doi 10.1016/j.cell.2017.05.035.
8. Muller S, Cho A, Liu SJ, Lim DA, Diaz A. CONICS integrates scRNA-seq with DNA sequencing to map gene expression to tumor sub-clones. Bioinformatics **2018**;34(18):3217-9 doi 10.1093/bioinformatics/bty316.
9. Alquicira-Hernández J, Sathe A, Ji H, Nguyen Q, Powell JE. scPred: Cell type prediction at single-cell resolution. bioRxiv2018.
10. Vento-Tormo R, Efremova M, Botting RA, Turco MY, Vento-Tormo M, Meyer KB*, et al.* Single-cell reconstruction of the early maternal-fetal interface in humans. Nature **2018**;563(7731):347-53 doi 10.1038/s41586-018-0698-6.

#### Legends to supplementary figures and tables

**Supplementary table S1**: Sequencing metrics, information on experimental batches and higher UMI count filter of all samples.  Rep 1 and 2 indicate technical replicates. Samples were prepared for 10X sequencing immediately after dissociation or after freeze-thawing (*).

**Supplementary table S2**: Marker genes used to define a cell lineage with literature reference.

**Supplementary table S3:** Cluster numbers filtered following reclustering analysis of respective cell lineages.

**Supplementary table S4:** Summary of clinical features of patient cohort used in the study. MSI = microsatellite instability, MSS = microsatellite stable, NA= not available

**Supplementary table S5:** Differentially expressed genes in epithelial cells of tumor, tumor-like and normal epithelium classes.

**Supplementary table S6:** Differentially expressed genes in respective macrophage clusters.

**Supplementary table S7:** Differentially expressed genes in respective DC clusters.

**Supplementary table S8:** Top 20 differentially enriched C7 MsigDB immunologic signatures (ANOVA FDR p value < 0.05) for respective myeloid lineage cluster.

**Supplementary table S9:** Top 20 differentially enriched regulons (ANOVA FDR p value < 0.05) for respective myeloid lineage cluster. Number in parenthesis indicates number of genes in the gene regulatory network.

**Supplementary table S10:** Differentially expressed genes in respective cytotoxic T cell clusters.

**Supplementary table S11:** Top 20 differentially enriched regulons (ANOVA FDR p value < 0.05) for respective lymphocyte lineage cluster. Number in parenthesis indicates number of genes in the gene regulatory network.

**Supplementary table S12:** Comparison of differentially expressed genes in tumor or normal stromal cells, with their logFC change and FDR p value, to the genes of the matrisome program.

**Supplementary table S13:** Top 20 differentially enriched regulons (ANOVA FDR p value < 0.05) for respective stromal cell lineage in tumor or normal tissue. Number in parenthesis indicates number of genes in the gene regulatory network.

**Supplementary table S14:** Tumor specific interactome with the identified receptor-ligand interaction and the two interacting cell types.

**Supplementary figure 1:** (A) Representative images of hematoxylin and eosin staining of FFPE tissue from respective patients. Scale bar indicates 50 μm. (B) UMAP representation of dimensionally reduced aggregated data from all samples following graph-based clustering with arbitrary cluster numbers (C) Dot plot depicting expression levels of specific genes across clusters with marker-based lineage assignments.

**Supplementary figure 2:** (A) UMAP representation for graph-based clustering of epithelial cells with arbitrary cluster numbers. (B) Expression of respective genes with grey indicating low and blue indicating high. (C) Heatmap depicting number of cells identified for each epithelial cell cluster by patient. (C) Expression of respective genes with grey indicating low and blue indicating high. (D) Distribution of posterior probabilities for cells to belong to the normal class or be unassigned following scPred analysis. (E) Heatmap depicting average gene set activity of top MSigDB Hallmark gene signatures following GSVA (ANOVA FDR p value < 0.05) across classes for respective patients.

**Supplementary figure 3:** (A) Heatmap representation of statistically significant copy number changes for depicted chromosomes from epithelial cells of respective patients. ‘amp’ denotes amplification, ‘del’ denotes deletion.

**Supplementary figure 4:** Heatmaps depicting average gene set activity of top MSigDB oncogenic c6 gene signatures following GSVA (ANOVA FDR p value < 0.05) across tumor epithelial clusters (containing at least 10 cells) for respective patients.

**Supplementary figure 5:** (A) Heatmap depicting number of cells identified for each cluster per patient. (B) Heatmap depicting average gene set activity of top MSigDB hallmark gene signatures following GSVA (ANOVA FDR p value < 0.05) across macrophage clusters. (C) Heatmap representation of significantly differentially expressed genes across DC clusters (log FC>0.25, FDR p value < 0.05)

**Supplementary figure 6:** (A) UMAP representation of lymphocytes following graph-based clustering with arbitrary cluster numbers and (B) cell lineages of each cluster. (C) Heatmap depicting expression of respective genes from each lymphocyte cluster. (D) Scatter plots depicting expression of respective genes in clusters 6 and 22 lymphocytes to allow lineage-based 'gating'.

**Supplementary figure 7:** (A) Heatmap depicting number of cells identified for each CD8 cluster by sample origin indicated by patient ID, normal (n) or tumor (t). 1 and 2 indicate replicates. (B) Heatmap depicting average gene set activity from MSigDB c7 GSE9650 across signatures cytotoxic T cell clusters(ANOVA FDR p value < 0.05) (C) Violin plots depicting expression of respective genes in c6 and c22 exhausted T cell clusters.

**Supplementary figure 8:** (A) Staining intensities for each single fluorescence channel per cell from five different regions of interest (ROI) from respective patient's tumor image. (B) Percentage of cells from five different regions of interest (ROI) classified as single-channel positive, multi-channel positive or negative for all channels from each patient's tumor image.

**Supplementary figure 9:** (A) Heatmap depicting number of Treg cells identified for each cluster according the sample origin. (B) Box plots depicting proportion of Tregs from total cells derived from tumor or normal site with z-test p value. (C) Heatmap depicting expression of respective genes for Treg clusters. (D) Heatmap depicting number of cells identified for each CD4 cluster by sample origin. (E) Violin plots depicting expression of respective genes in CD4 T cells.

**Supplementary figure 10:** (A) Heatmap depicting number of cells identified for each NK cell cluster by sample origin. (B-C) Heatmap depicting expression of respective genes from each NK cell cluster. (D) Heatmap depicting number of cells identified for each B cell cluster by sample origin. (E) Heatmap depicting number of cells identified for each plasma cell cluster by sample origin. (F) Violin plots depicting expression of respective genes in plasma cells.

**Supplementary figure 11:** Heatmaps depicting number of cells identified in (A) fibroblast and (B) endothelial cell lineage in each patient. (C) Heatmap representation of significantly differentially expressed genes across endothelial cell clusters (log FC>0.25, FDR p value < 0.05)  (D) Violin plot depicting expression of *ACTA2* with FDR p value.

#### Supplementary figure 1

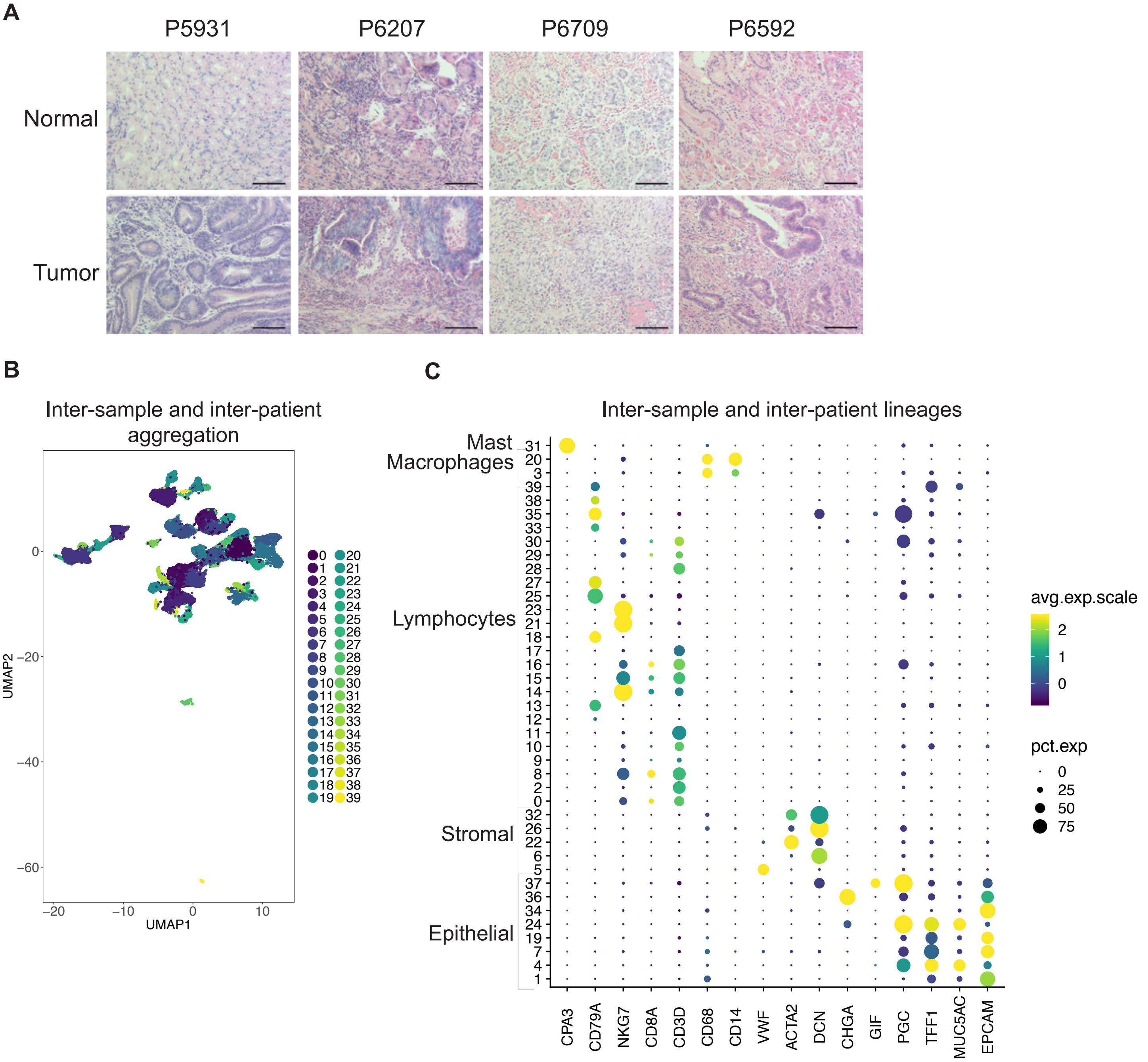

#### Supplementary figure 2

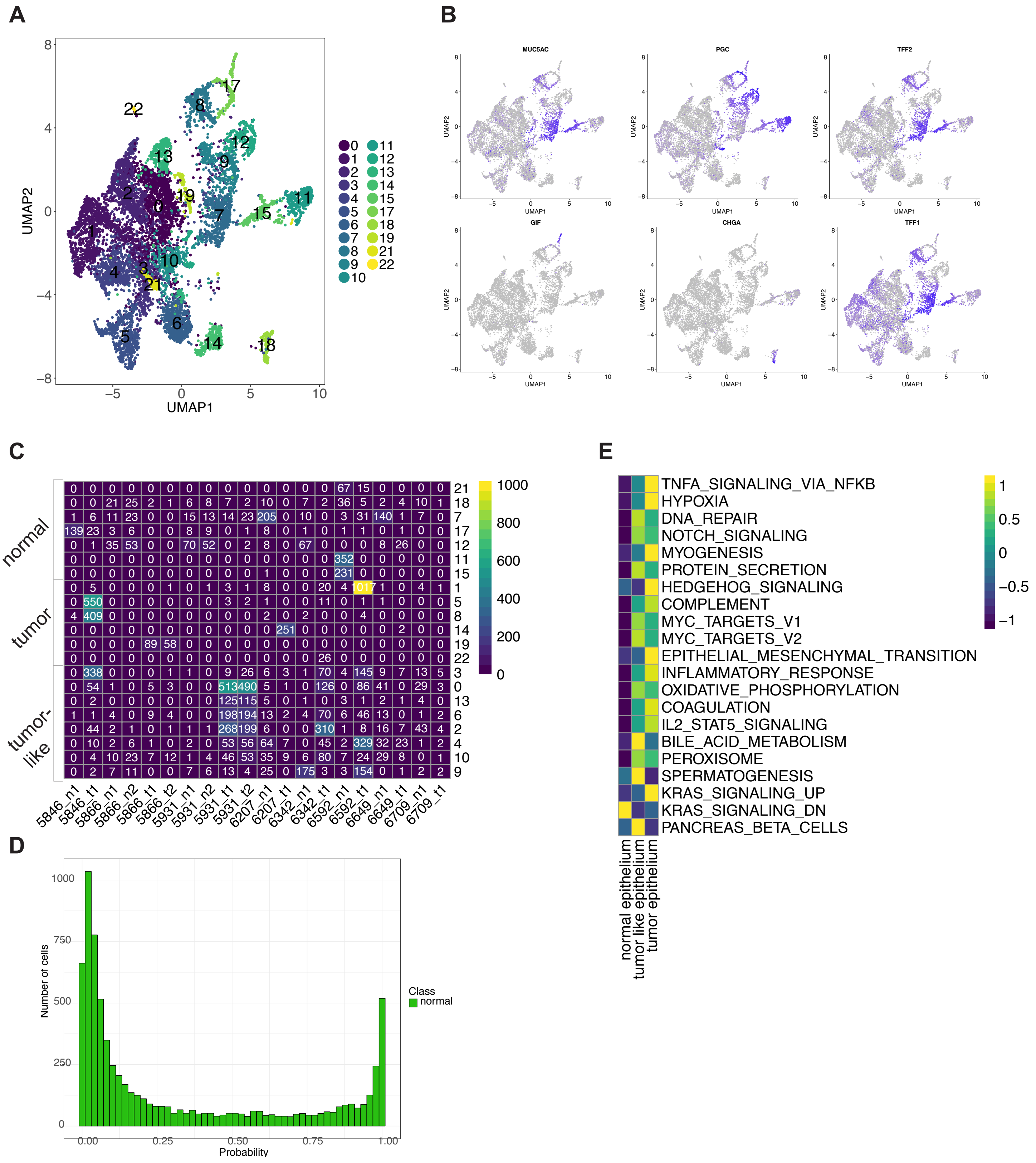

### Supplementary figure 3

**A**

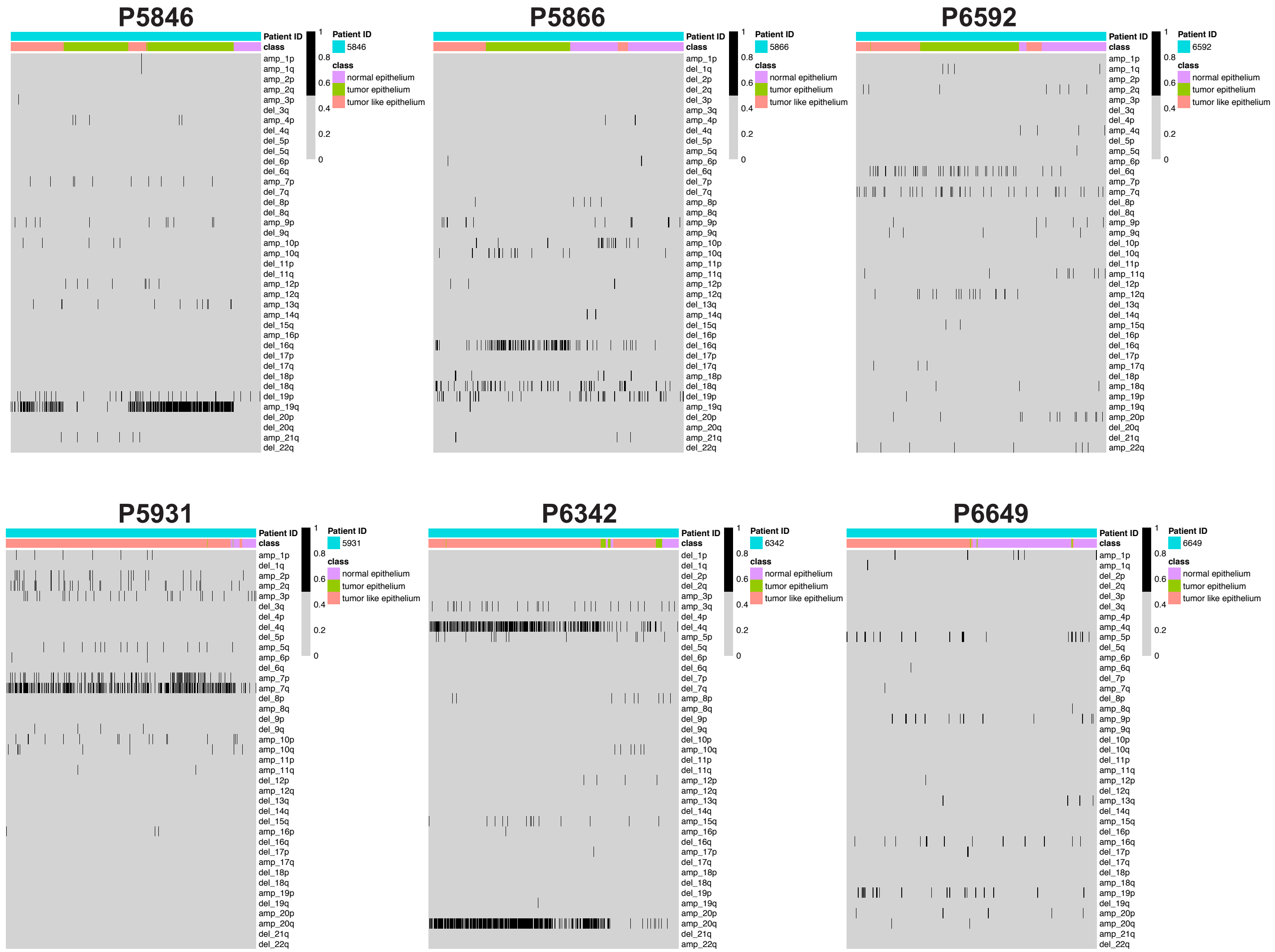

#### Supplementary figure 4

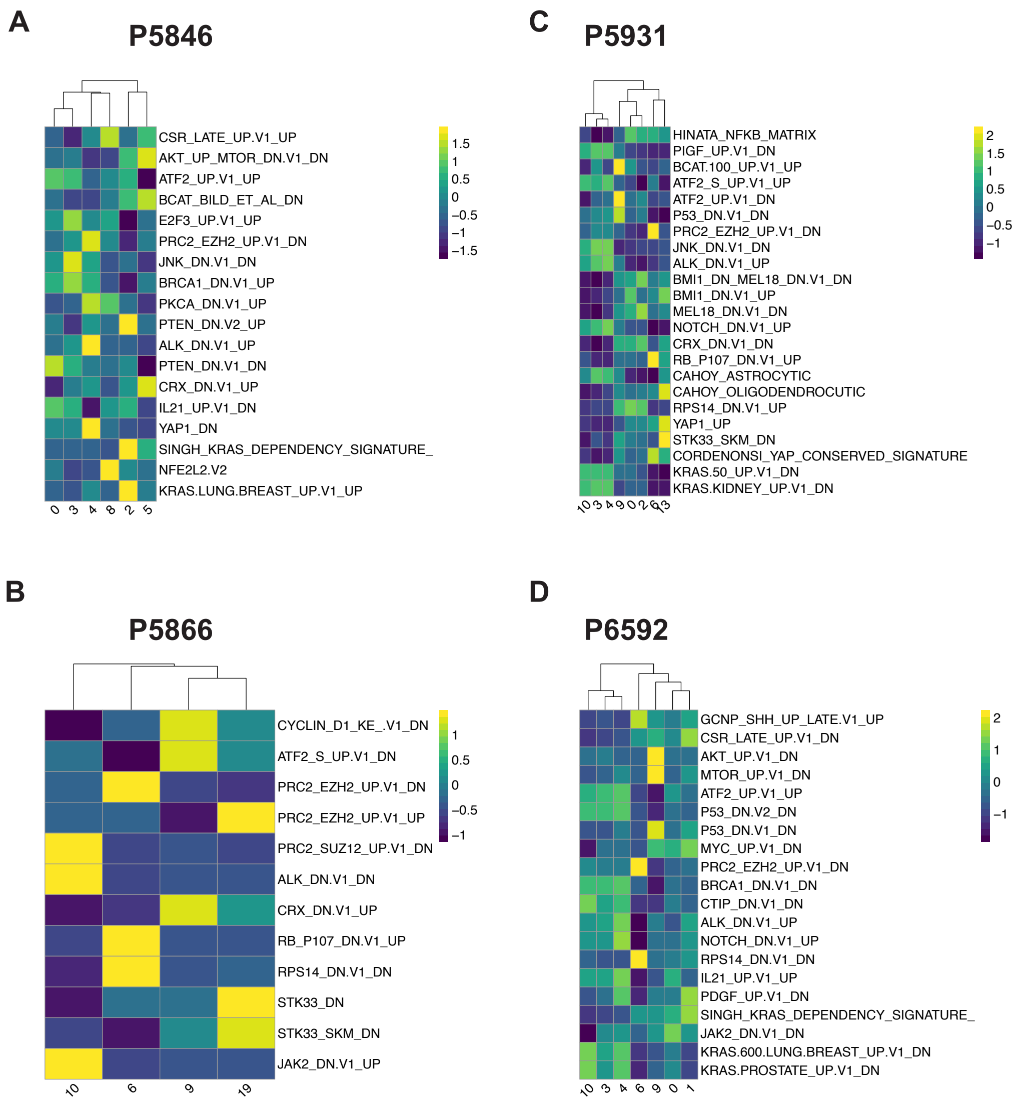

#### Supplementary figure 5

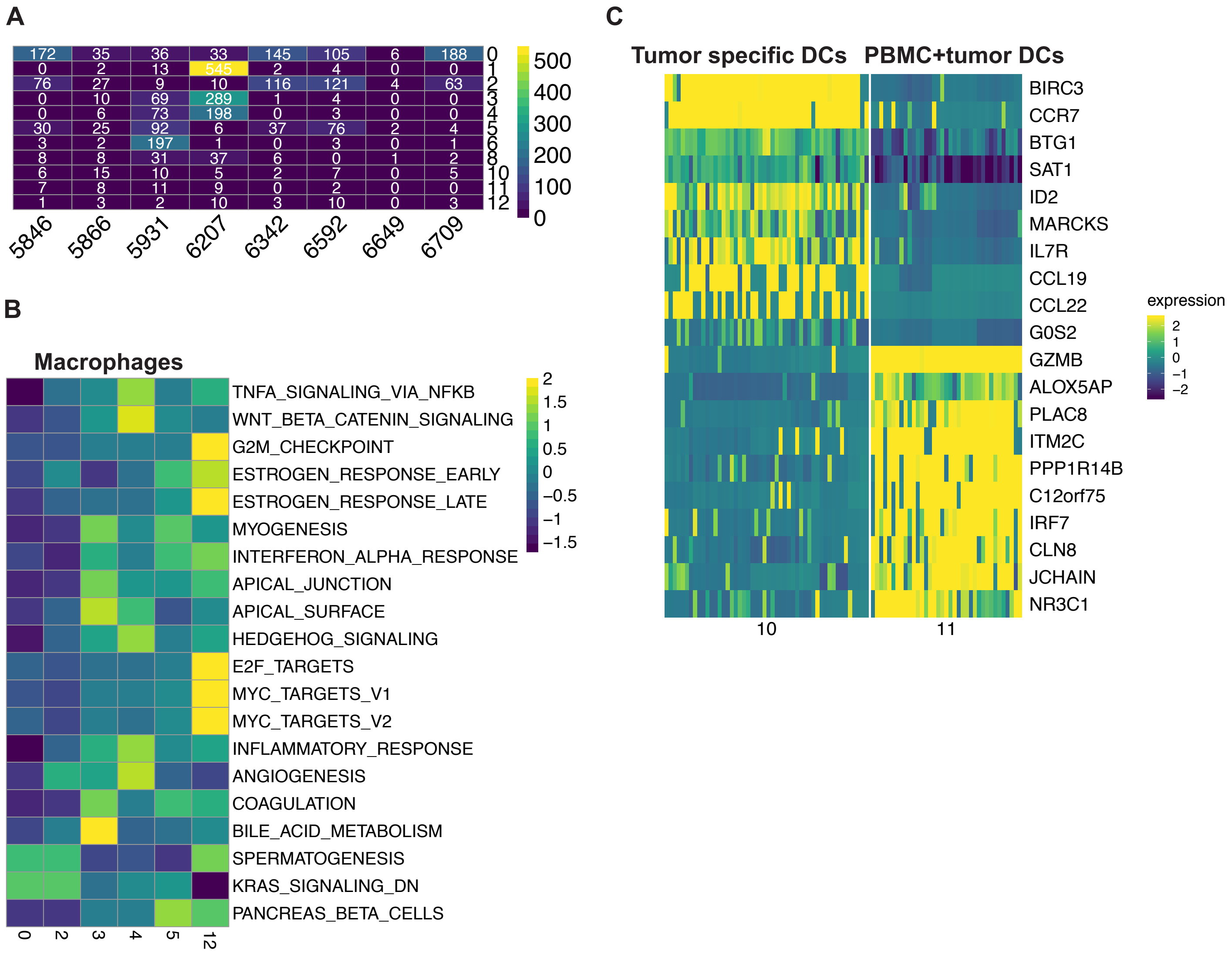

### Supplementary figure 6

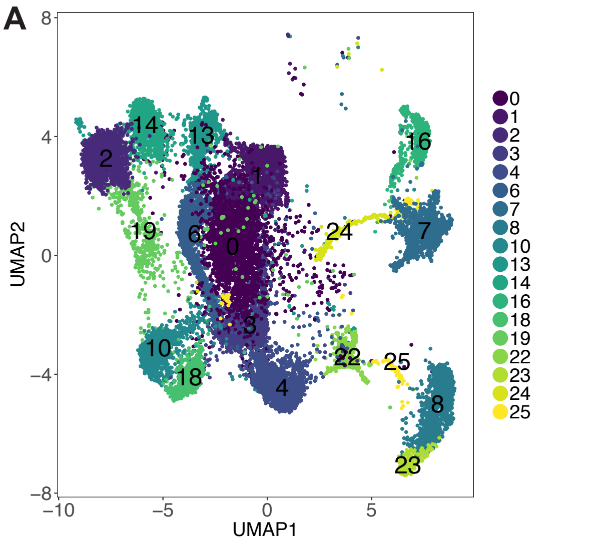


B

| Cell type | Cluster number |
|---|---|
| CD8 | 0,1,6,19,22 |
| CD4 | 3,6,10,18 |
| NK | 2,13,14 |
| B | 8,22,23 |
| Plasma | 7,16,24,25 |
| Treg | 4,22 |

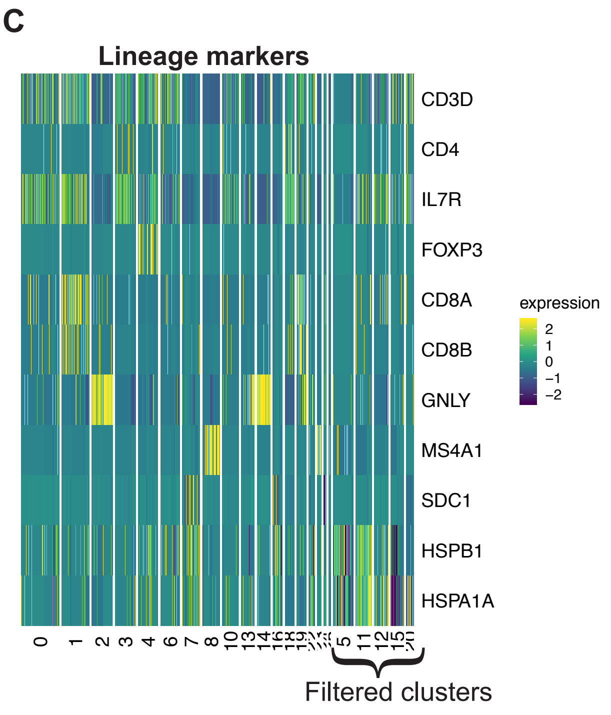


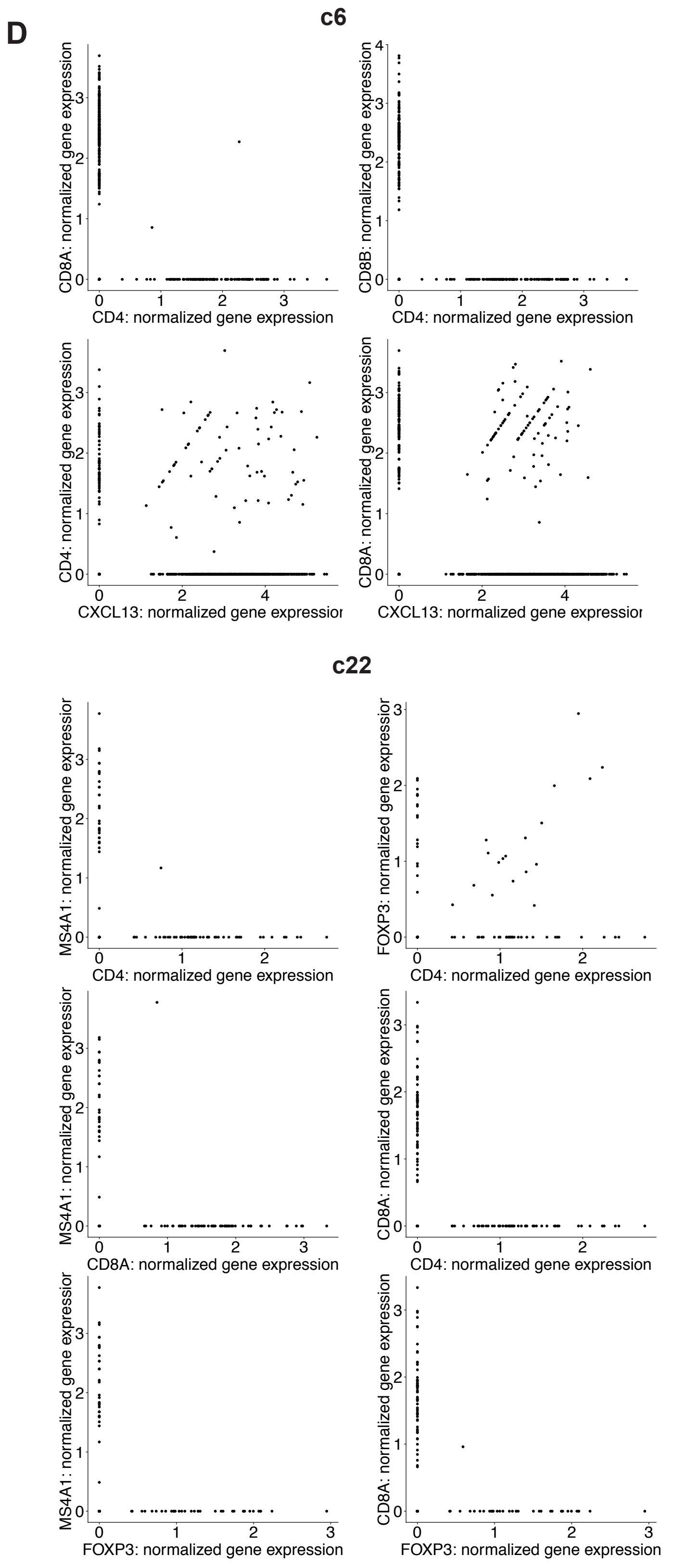

#### Supplementary figure 7

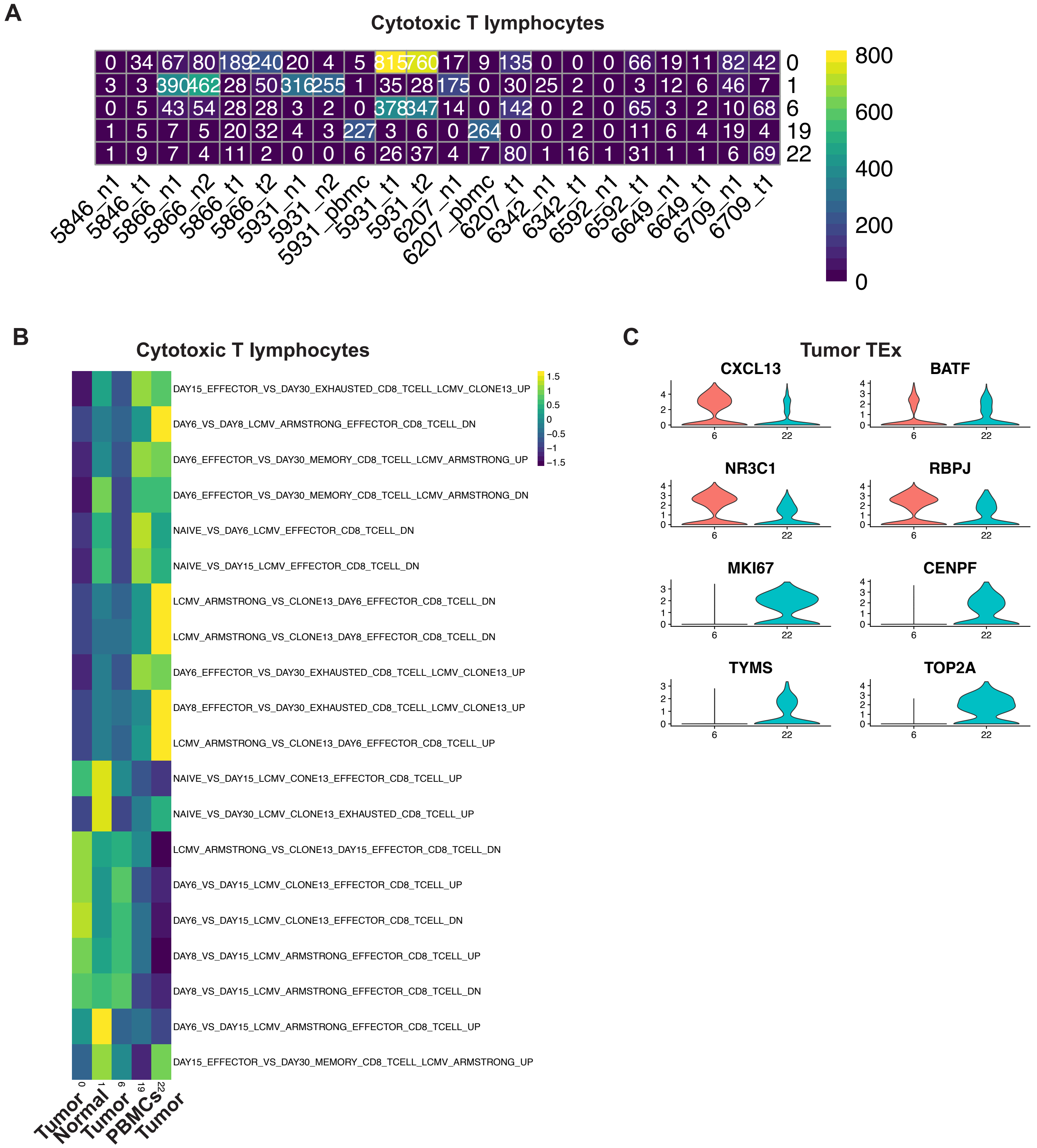

### Supplementary figure 8

## A

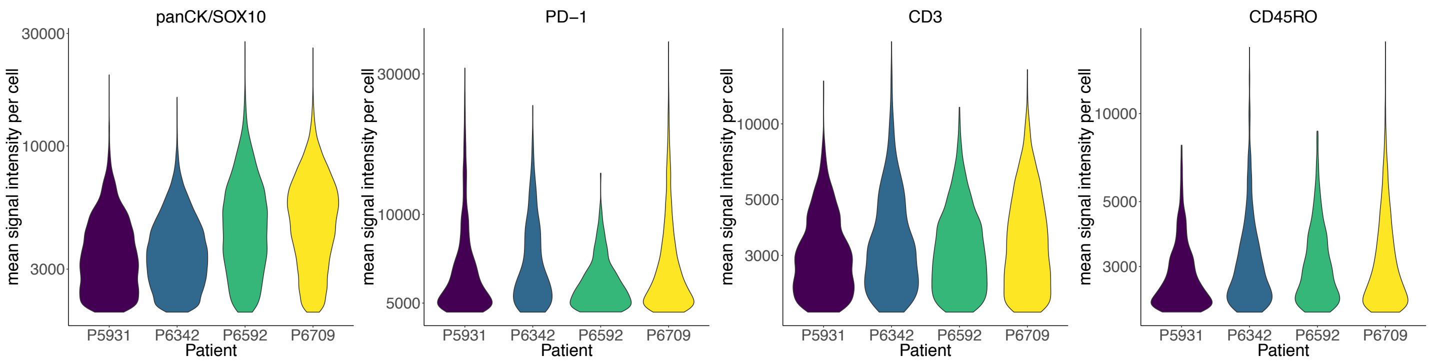


## B

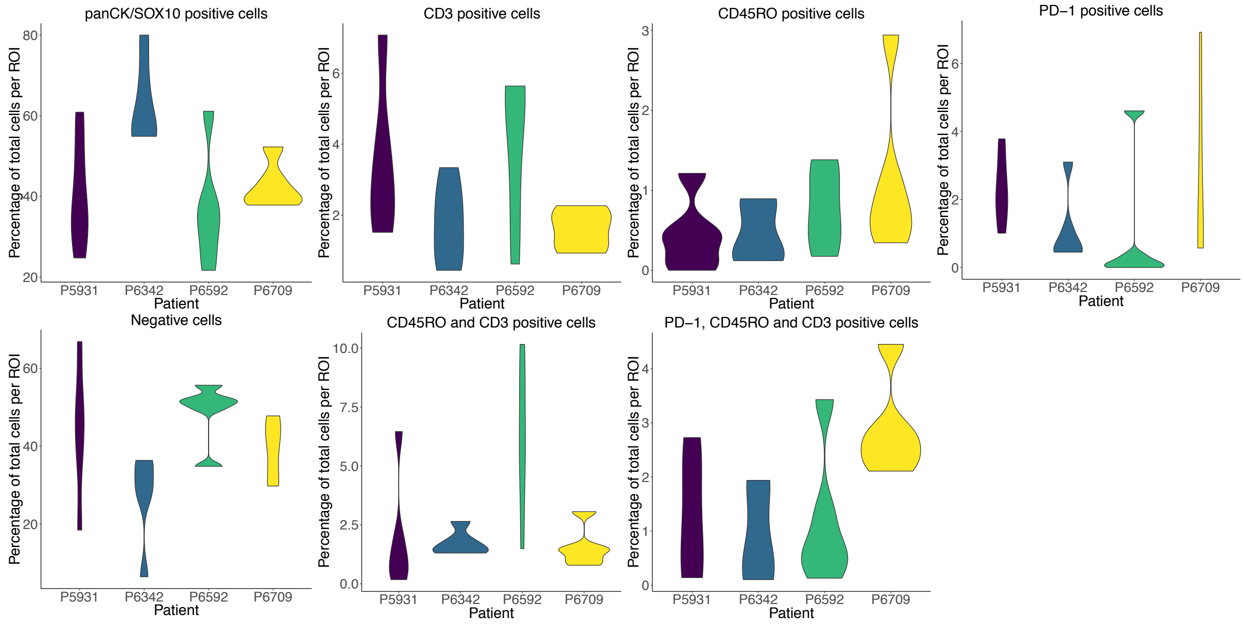

### Supplementary figure 9

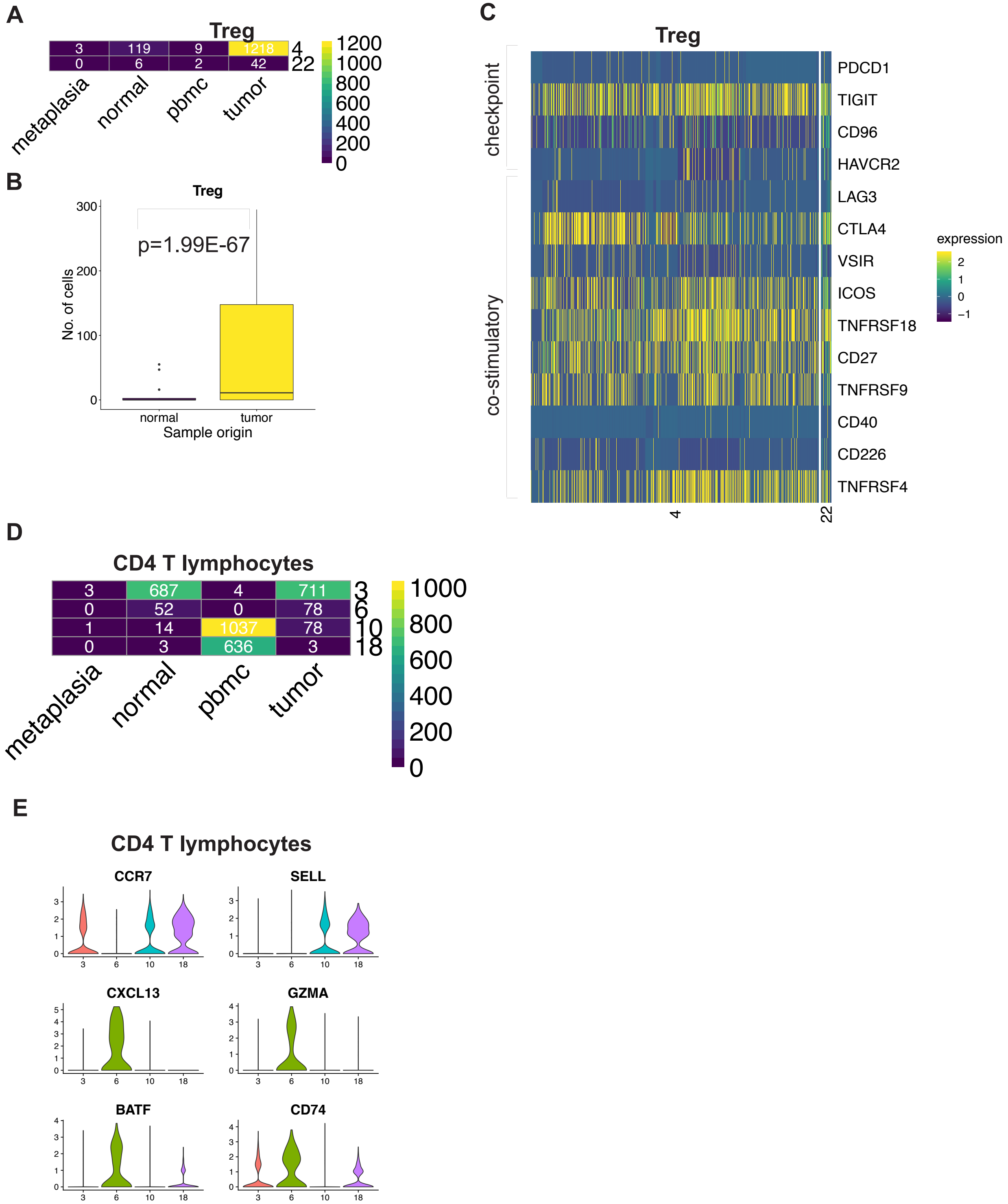

#### Supplementary figure 10

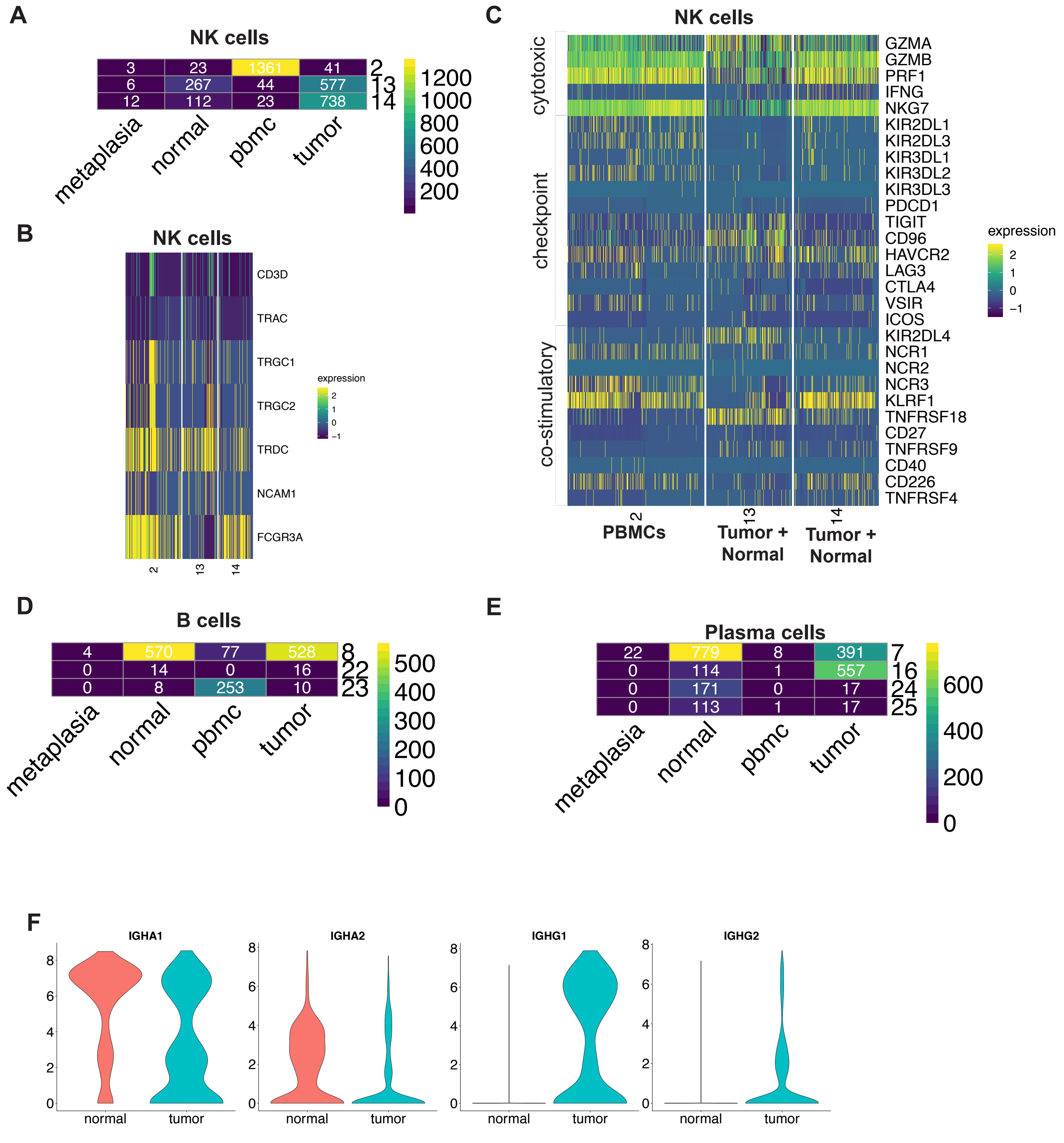

### Supplementary figure 11

**A**

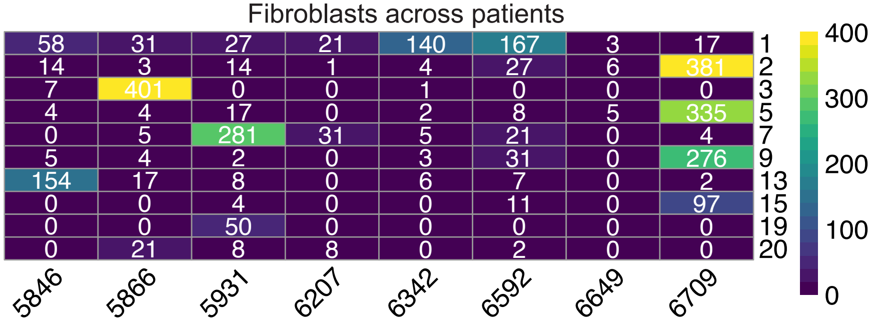


**B**

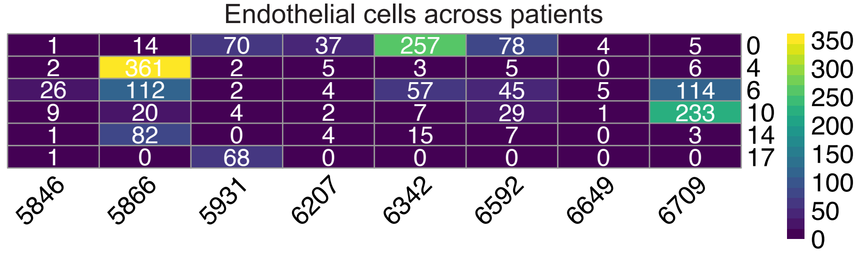


**C**

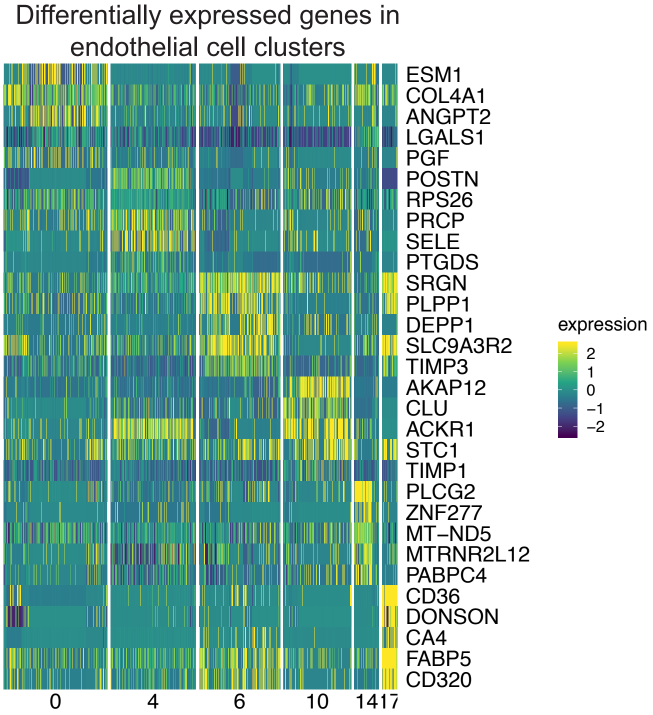


**D**

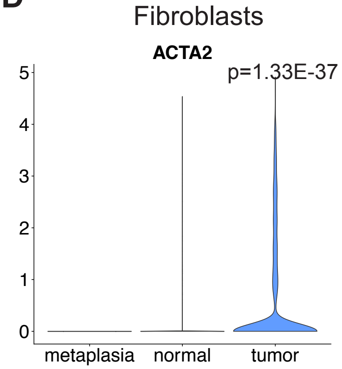
